# A general role for ventral white matter pathways in morphological processing: going beyond reading

**DOI:** 10.1101/2020.06.21.163303

**Authors:** Maya Yablonski, Benjamin Menashe, Michal Ben-Shachar

## Abstract

The ability to recognize the structural components of words, known as morphological processing, was recently associated with the bilateral ventral white matter pathways, across different writing systems. However, it remains unclear whether these associations are specific to the context of reading. To shed light on this question, in the current study we investigated whether the ventral pathways are associated with morphological processing in an oral word production task that does not involve reading. Forty-five participants completed a morpheme-based fluency task in Hebrew, as well as diffusion MRI (dMRI) scans. We used probabilistic tractography to segment the major ventral and dorsal white matter pathways, and assessed the correlations between their microstructural properties and performance on the morpheme-based fluency task. We found significant correlations between morpheme-based fluency and properties of the bilateral ventral tracts, suggesting that the involvement of these tracts in morphological processing extends beyond the reading modality. In addition, significant correlations were found in the frontal aslant tract (FAT), a dorsal tract associated with oral fluency and speech production. Together, our findings emphasize that neurocognitive associations reflect both the cognitive construct under investigation as well as the task used for its assessment. Lastly, to elucidate the biological factors underlying these correlations, we incorporated the composite hindered and restricted model of diffusion (CHARMED) framework, measured in independent scans. We found that only some of our findings could be attributed to variation in a CHARMED-based estimate of fiber density. Further, we were able to uncover additional correlations that could not be detected using traditional dMRI indices. In sum, our results show that the involvement of the ventral tracts in morphological processing extends to the production domain, and demonstrate the added value of including sensitive structural measurements in neurocognitive investigations.

## 1. Introduction

A central goal in cognitive neuroscience is understanding the language circuitry in the brain. Numerous neuroimaging studies have revealed that cortical regions along the left dorsal route are involved in phonological functions (mapping speech sounds to articulatory programs), while ventral regions are involved in comprehension and meaning extraction (Hickok and Poeppel, 2007; Price, 2012; Saur et al., 2008). In addition, studies focusing on reading processes consistently find that regions along the ventral occipito-temporal cortex are involved in visual processing of written language (Cohen et al., 2000; Lerma-Usabiaga et al., 2018; McCandliss et al., 2003; Vinckier et al., 2007; White et al., 2019). Since language processing requires integrating different types of information at multiple levels, it is imperative to include in this account the underlying structural connectivity that enables representational transformations along these cortical pathways. Indeed, structural properties of white matter tracts were found to be associated with specific language skills, complementing and expanding functional findings. Dorsal tracts in the left hemisphere were shown to predict phonological skills (Saygin et al., 2013; Yeatman et al., 2011), while ventral tracts were broadly associated with access to meaning (Nugiel et al., 2016; Sarubbo et al., 2015) as well as with reading ability (Cummine et al., 2015; Ozernov-Palchik et al., 2019; Vandermosten et al., 2012; Yeatman et al., 2012a). However, it remains an open question whether these white matter tracts can be mapped onto specific language processes across stimulus and response modality, such as reading, listening and speaking.

One of the linguistic functions that has been relatively understudied in relation to white matter is morphological processing. Morphological processing refers to the ability to decompose complex words into their structural units, morphemes. For example, a complex word, such as ‘endlessly’, can be broken down to three constituent morphemes: ‘end’+’less’+’ly’. Morphological awareness, the ability to recognize and manipulate morphemes, is a strong predictor of reading ability, both in children (Kirby et al., 2012; Nagy et al., 2006; Ravid and Mashraki, 2007; Saiegh-Haddad and Taha, 2017; Schiff et al., 2011; Siegel, 2008) and in adults (Law et al., 2015; Metsala et al., 2019; Tighe and Binder, 2015). This is true even when controlling for other predictive factors such as phonological awareness, the ability to identify and manipulate speech sounds. Poor readers often demonstrate impaired morphological knowledge (Law et al., 2015; Schiff and Raveh, 2007; Schiff and Ravid, 2013, 2007; Tong et al., 2014) and benefit from intervention programs that specifically target morphological skills (Arnbak and Elbro, 2000; Bowers et al., 2010; Goodwin and Ahn, 2010; Taha and Saiegh-Haddad, 2015). Despite the breadth of behavioral evidence emphasizing the invaluable role of morphological processing in reading, its place within the broader language “connectome” in the brain has been generally overlooked (Catani and Bambini, 2014; Dick et al., 2014; Dick and Tremblay, 2012; Friederici and Gierhan, 2013).

Recently, we have demonstrated in two independent samples of English and Hebrew readers, an association between sensitivity to the morphological structure of written words and microstructural properties of the bilateral ventral tracts (Yablonski et al., 2019; Yablonski and Ben-Shachar, 2020). We interpreted these associations as suggesting that morphological information plays a role in lexical access, a process in which readers activate the mental representations of words, supported by the bilateral ventral pathways. This is in line with accounts that view morphemes as a key factor in mapping between written word forms and their meanings (Rastle, 2019). The associations between morphological processing and the ventral tracts emerged in both English and Hebrew, languages that greatly differ in their structure. This neurobiological generalization across orthographies and morphological systems supports a notion of morphological processing as a cognitive process that abstracts away from the specifics of the writing system.

Critically, the ventral tracts have previously been implicated in visual processing of written words without any morphological manipulations (Cummine et al., 2015; Huber et al., 2018; Welcome and Joanisse, 2014; Yeatman et al., 2012a). Thus, an alternative explanation of the correlations observed between the ventral tracts and morphological processing may suggest that they arise from task-specific demands, imposed by the visual presentation of written stimuli in our previous studies. In other words, it remains unclear whether the involvement of the ventral tracts in morphological processing is specific to the visual modality of reading, or generalizes across input modalities. To shed light on this question, the aim of the current study was to investigate whether the ventral tracts are associated with morphological processing in settings that do not involve reading. Further, we sought to explore whether the involvement of the ventral tracts in morphological processing extends beyond perception to the domain of language production.

To this end, we used a morpheme-based fluency task, where participants were required to generate as many words as possible that share a morpheme with the target word. For example, upon hearing the word ‘likely’, a participant could respond with ‘likeable’, ‘unlike’, ‘likelihood’ etc., which all share the same base morpheme, the stem. Alternatively, participants could be asked to generate words that share a suffix with the target, to which they could respond with ‘wildly’, ‘quickly’, ‘partly’ etc. Target words were read aloud by the experimenter, and participants produced their responses orally. This task thus requires the participant to first extract the target morpheme (stem or affix) from the word they heard, then search their mental lexicon for words that share the same morpheme, and say them out loud. Although very few studies have used this task, there are several reports that dyslexics generate less morphologically-related words compared with typical readers (Ben-Dror et al., 1995; Leikin and Even Zur, 2006; Su et al., 2018; but see Casalis et al., 2004). Importantly, the task does not involve any reading material or visual presentation of stimuli. In this manner, this task taps into morphological processing without explicit orthographic processing. Here, participants performed the morpheme-based fluency task in Hebrew and completed diffusion MRI scans, which allowed us to reconstruct their dorsal and ventral white matter tracts of interest. We then inspected the neurocognitive correlations between their morpheme-based fluency and microstructural properties of their white matter tracts.

An additional goal of the current study was to improve our understanding of the white matter tissue properties that underlie morphological sensitivity. A major challenge in diffusion MRI studies is interpreting the underlying tissue components that give rise to the measured diffusion signal. Commonly used diffusion-based metrics, such as fractional anisotropy (FA) and mean diffusivity (MD), are very useful in detecting correlations with behavioral measurements but prove difficult to interpret, because their relation to tissue properties is indirect and ambiguous. For example, FA is affected by myelin, axonal density, axonal diameter and directional coherence, so FA correlations may be driven by each of these factors or by their interactions. Several models have been proposed to separately estimate the diffusion signal that originates from different cellular compartments (Alexander et al., 2010; Jensen et al., 2005; Tuch et al., 2002). Here, we estimated the composite hindered and restricted model of diffusion (CHARMED; Assaf et al., 2004; Assaf and Basser, 2005), which separates the signal decay into hindered (extracellular) and restricted (intracellular) compartments. The CHARMED framework requires acquiring multi-shell diffusion data at several b-values, and is used to calculate, within each voxel, the volume fraction of the restricted compartment (FR). Indeed, FR was shown to be more sensitive to structural changes in both gray and white matter compared to tensor-derived metrics (De Santis et al., 2017; Tavor et al., 2013). We therefore acquired a full CHARMED dataset in our participants and incorporated FR as an additional measure of interest in the current study.

In all, 45 healthy adult Hebrew readers completed the morpheme-based fluency task and underwent diffusion MRI (dMRI) scans in separate behavioral- and MRI-sessions. We used constrained spherical deconvolution (CSD) to model diffusivity at the voxel level, followed by probabilistic fiber tractography to identify the major ventral and dorsal language tracts in each participant. In a parallel processing stream we applied tensor modelling and deterministic tractography to estimate the same pathways. We then evaluated the correlations between morpheme-based fluency scores and microstructural properties along the ventral white matter tracts of interest. Based on our previous findings, we targeted the inferior fronto-occipital fasciculus (IFOF), inferior longitudinal fasciculus (ILF) and uncinate fasciculus (UF), bilaterally. In addition, we included dorsal tracts that are known to be involved in speech production, namely, the fronto-temporal and fronto-parietal segments of the arcuate fasciculus (AF-ft, AF-fp, respectively), and the frontal aslant tract (FAT) (Catani et al., 2013; Fridriksson et al., 2013; Kronfeld-Duenias et al., 2016; Marchina et al., 2011). We assessed the specificity of the identified associations using multiple regression models, controlling for performance on standard verbal fluency tasks that do not depend on morphological processing in any explicit way. Specifically, we administered the widely used letter-based and category-based fluency tasks, where participants generate words that begin with an opening letter or belong to a given semantic category (e.g., animals). Lastly, we investigated correlations between morpheme-based fluency and FR, first as a follow-up analysis within clusters showing significant FA or MD associations, and later as an exploratory analysis applied to all tracts of interest.

Our hypotheses followed this line of reasoning: On the one hand, prior associations between morphological processing and the ventral tracts may reflect a general role for the ventral tracts in decomposing a word into its morphemes and accessing its lexical representation. If this is true, then the ventral tracts are expected to be associated with different tasks that rely on morpheme-mediated lexical access, regardless of presentation and response modality. In this case, we expect to find correlations between morpheme-based fluency and properties of the bilateral ventral tracts, as in our previous studies. On the other hand, previously observed effects may have reflected a task-specific aspect of morphological processing, related specifically to its implementation in visuo-orthographic processing. In this case, the ventral tracts are not expected to be involved in morphological processing in other modalities. Instead, morpheme-based fluency may map onto dorsal tracts, previously associated with other measures of verbal fluency (Blecher et al., 2019; Catani et al., 2013). Testing these hypotheses would lead to better understanding of the involvement of ventral and dorsal language tracts in morphological processing across task and modality.

## 2. Methods

### 2.1. Participants

Forty-five Hebrew speaking adults participated in this study (mean age 26.45 ± 3.72 years, age range 20-35, 16 males). Participants were recruited as part of a larger research project (Yablonski and Ben-Shachar, 2020) which included extensive behavioral assessment and MRI scans, conducted in separate sessions. All participants were right handed, as estimated by the Edinburgh handedness inventory (Oldfield, 1971; see Supplementary Table S1), had normal hearing and vision, and no known history of diagnosed learning disabilities or neurological conditions. Only participants whose native language is Hebrew were included in the study. One participant was referred to neurological follow-up due to an incidental finding, but otherwise had normal anatomical structure and diffusion values and was thus not excluded from analysis. Participants were paid 200 NIS for their participation. The research was approved by the Helsinki Committee of Sheba Medical Center, by the Institutional Review Board of Tel Aviv University, and by the Ethics committee of the Faculty of Humanities in Bar-Ilan University. All participants signed a written informed consent before participating in the study.

### 2.2. Cognitive assessment

Participants completed an extensive cognitive assessment battery which included several measures of reading related skills, reported in a previous publication (Yablonski and Ben-Shachar, 2020). The cognitive assessment took place in a quiet room, a couple of weeks before the MRI scan. In the current analysis we focused on the verbal fluency tasks, which were not included in the previous analysis (for fMRI results pertaining to the fluency task see Agmon et al., submitted).

#### 2.2.1. Morpheme-based fluency

To better understand this task, we first briefly introduce the Hebrew morphological system. The majority of Hebrew words comprise two interleaved morphemes, such that a tri-consonantal *root* morpheme is embedded within a phonological template, the *pattern* morpheme. When a root is combined with different patterns, it results in a family of derived words, sharing the 3 root consonants and often related in meaning. For example, the root *SGR* can be embedded in various patterns to generate *SaGaR* (he closed; root letters are denoted by capital letters), *miSGeRet* (a frame) or *heSGeR* (quarantine). Additional affixes may be added to these non-linear combinations to inflect the word for different grammatical features, such as gender, number, or tense (e.g., *SaGaR* can be inflected to generate *SaGaRti*, I closed, or *SaGaRnu*, we closed). Hebrew speakers are able to utilize root and pattern information as early as the age of three (Clark and Berman, 1984), and this ability gradually improves throughout the school years (Bar-on and Ravid, 2011; Ravid and Malenky, 2001; Ravid and Schiff, 2006). The systematic nature in which roots and patterns combine to form words has led to the view that the Hebrew mental lexicon is organized around word-families, clusters of words that share the same root or pattern morpheme (Deutsch et al., 1998; Deutsch and Meir, 2011; Frost, 2012; Frost et al., 1997; Ravid and Schiff, 2006).

In the morpheme-based fluency tasks used here, we used Hebrew morphemes as cues to initiate lexical search. Participants were asked to generate as many words as possible that shared a morpheme (either a root or a pattern) with a spoken target word. For the root-based fluency task, the instruction was to come up with as many words as possible that shared a root with the target. For example, upon hearing the target word *tiZMoRet*, an orchestra, participants would generate words like *ZaMeRet*, a female singer, *ZaMiR*, a nightingale, and so on, all derived from the same root morpheme, *ZMR* (see illustration in Figure 1). For the pattern-based fluency task, participants had to come up with as many words as possible that shared a pattern with the target. For example, upon hearing the word *maMTeRa*, a sprinkler, participants generated *maDReGa*, a stair, *maVReGa*, a screwdriver, etc.

**Figure 1.**
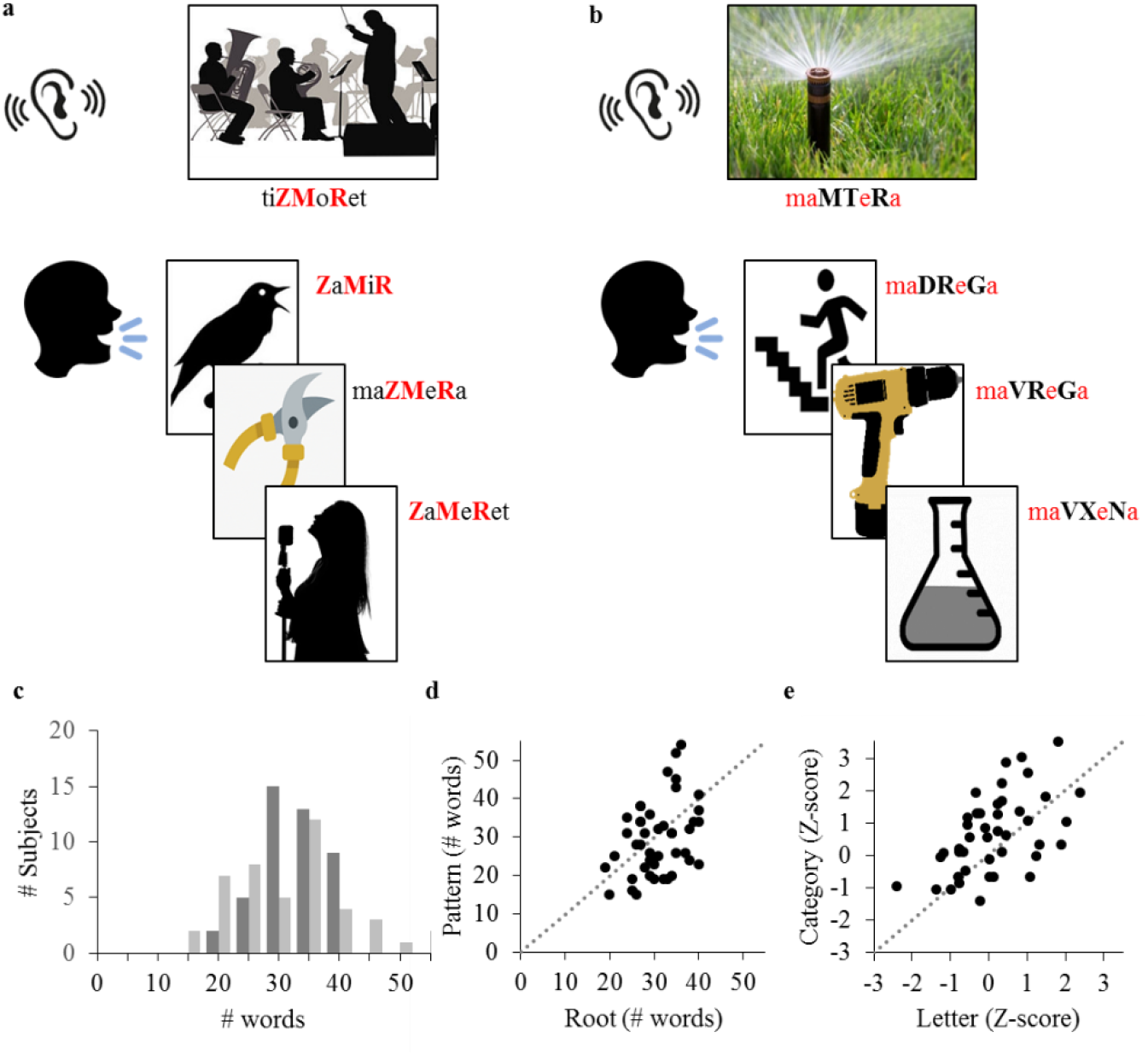
Morpheme-based fluency tasks. In the root-based fluency task (a), participants are required to generate as many words as possible that share a root with the target, but have different patterns. In the pattern-based fluency task (b), participants are required to generate as many words as possible that share a pattern with the target but have different roots. Upon hearing the target word, participants produced word responses orally. In both panels, root consonants appear in uppercase, while the shared morpheme (root or pattern) in each task is colored in red. Phonetic transcripts appear next to each word to emphasize the morphological structure and overlap between target and responses, no written stimuli were included in the actual task. Performance in the two tasks was variable across individuals (c; Dark gray-root, light gray-pattern) and weakly correlated (d; r = 0.38, p < 0.05 uncorrected). Performance in the two control tasks, letter-based fluency and category-based fluency, was highly correlated (e; r = 0.53, p < 0.05, FDR corrected, see Supplementary Table S3 for the full correlation matrix of the behavioral tasks). Dashed lines delineate y = x.

Before each task, participants were given two example items to make sure they understood the instructions. Participants received feedback for their responses to the example items. In case they couldn’t come up with correct responses at first, the experimenter provided several example responses. The instructions put emphasis on the distinction between derivations (different words) and inflections (syntactic modulations of the same word; i.e., “We are interested in your ability to come up with different words that would appear as separate entries in a dictionary, not in different forms of the same word. For example, if you say “dance”, please don’t add “dancing, danced, will dance”). Each task was sampled using 5 different target items, and participants were given 30s to produce their responses to each target. The 5 items were selected out of 10 used in (Leikin and Even Zur, 2006), in order to keep the total duration of the cognitive battery around 1hr. See supplementary Table S2 for a list of target items used in each task.

#### 2.2.2. Letter-based and category-based fluency

The standard verbal fluency tasks were presented according to the normed Hebrew version described in (Kavé, 2005; Kavé and Knafo-Noam, 2015; see The Delis-Kaplan Executive Function System (Delis et al., 2001) for the original English version of this test). This tool is widely used in clinical assessment (Shao et al., 2014), encompassing a letter-based fluency task and a category-based fluency task. In both tasks, participants were provided with a criterion (a single opening letter or a semantic category), and were asked to provide as many Hebrew words as possible that satisfy this criterion within 60 s. Each task is repeated 3 times with different criteria. For letter-based fluency, participants were asked to produce words that begin with the letters Bet (/b/), Gimel (/g/), or Shin (/sh/ or /s/). For category-based fluency, participants were asked to produce words that belong to the categories animals, fruits and vegetables, and vehicles.

#### 2.2.3. Procedure

The administration order of tasks and of items within tasks was kept constant across participants: The letter-based fluency task was administered first, followed by the category-based fluency task, root-based fluency and pattern-based fluency tasks. The experimenter read each target aloud and participants’ oral responses were recorded and transcribed offline. Lastly, participants completed a timed single word reading test, where they were asked to read aloud as many words as possible in 1 minute, without compromising accuracy (Shatil, 1997).

#### 2.2.4. Fluency score analysis

As a first step, repetitions and erroneous responses were removed. In the morpheme-based tasks (root and pattern fluency), additional screening was applied to exclude responses that were inflections of the same word (e.g., playing, played). In such cases, only the first response was taken into account. The score on each task (root-based and pattern-based) was the total number of correct unique responses, summed across the 5 target items.

Responses in the letter-based and category-based tasks were screened according to the guidelines described in (Kavé and Knafo-Noam, 2015). For example, in the category-based task, names of subcategories (e.g., bird) were not counted if the participant also produced specific exemplars within that subcategory (e.g., raven, sparrow). The total number of correct unique responses was coded per criterion and summed across the three criteria within each task. Then, a standardized score was calculated based on age-appropriate Hebrew norms (Kavé and Knafo-Noam, 2015). Subjects whose score in any of the fluency tasks deviated by more than 2.5 standard deviations (SD) from the group mean were excluded as outliers from that particular task (but were kept in analyses of other tasks if their score was within the 2.5 SD range).

### 2.3. White matter analysis

#### 2.3.1. MRI data acquisition

MRI scans were conducted at the Strauss Center for Computational Neuroimaging at Tel Aviv University. Data were collected using a 3T Siemens Magnetom Prisma scanner (Siemens Medical Systems, Erlangen, Germany), with a 64-channel head coil. A standard dMRI protocol was applied by means of a single-shot spin-echo diffusion-weighted echo-planar imaging sequence (86 axial slices, each 1.7 mm thick, no gap; FOV=204 × 204 mm, image matrix size=120 × 120 providing a cubic resolution of 1.7 × 1.7 × 1.7 mm, TR = 4000 ms, TE = 58 ms). Sixty-four diffusion-weighted volumes (b=1000 s/mm^2^) and three reference volumes (b=0 s/mm^2^) were acquired using a standard diffusion direction matrix. Multiband acceleration was used with slice acceleration factor of 2. Total acquisition time for the dMRI sequence was 4:48 min.

Data for the CHARMED analysis were acquired using a spin-echo diffusion-weighted echo-planar imaging sequence with 64 axial slices, each 2.2 mm thick, and no gap. The scanning parameters were FOV=202 × 202 mm, image matrix size=92 × 92 providing a cubic resolution of 2.2 × 2.2 × 2.2 mm, TR = 3600 ms, TE = 70 ms, with diffusion pulse separation/duration of Δ/δ = 38/18 ms. The following gradient scheme was used to acquire multi-shell diffusion weighted data: 20 directions with b=250 s/mm^2^, 64 directions with b=1000 s/mm^2^, 30 directions with b=2500 s/mm^2^, 30 directions with b=4000 s/mm^2^, and 6 reference volumes (b=0 s/mm^2^). Multiband acceleration was used with slice acceleration factor of 2. Total acquisition time for the CHARMED sequence was 9:32 min.

In addition, high resolution T1-weighted anatomical images were acquired for each participant using a magnetization prepared rapid acquisition gradient echo (MPRAGE) protocol (TR = 2530 ms, TE = 2.99 ms, flip angle = 7°, 1 mm thick slices, matrix size: 224×224×176, voxel size: 1× 1×1mm). Lastly, participants also completed a functional MRI protocol which was analyzed and reported in a separate publication (Agmon et al., submitted).

#### 2.3.2. dMRI analysis

Throughout this paper, we report results obtained using the constrained spherical deconvolution (CSD) model at the voxel level and probabilistic tractography for tract reconstruction (Tournier et al., 2007). To allow comparison with our previous studies (Yablonski et al., 2019), we ran a parallel analysis pipeline using the diffusion tensor model at the voxel level and tract reconstruction with deterministic tractography (Basser et al., 2000; Mori et al., 1999). The results obtained using the two pipelines were quite similar. For simplicity, we report the CSD+probabilistic tractography results in the main text, and the tensor+deterministic tractography results in the Supplementary material. In general, our analysis procedure consisted of three steps, all conducted within the native space of each participant: (1) preprocessing of the raw diffusion data, including model fitting (tensor and CSD) at the voxel level (2) whole brain tractography and (3) automated tract segmentation and quantification.

##### 2.3.2.1. Preprocessing

Initial preprocessing was implemented using ‘mrDiffusion’ (https://github.com/vistalab/vistasoft), an open-source package implemented in Matlab 2012b (The Mathworks, Nattick, MA). T1 images were aligned to the anterior commissure - posterior commissure (AC-PC) orientation. Diffusion weighted images were corrected for eddy-current distortions and head motion (Rohde et al., 2004). Diffusion-weighted volumes were registered to the averaged non-diffusion weighted (b0) volume, which was registered to the T1 image using a rigid body mutual information maximization algorithm (implemented in SPM8; Friston and Ashburner, 2004). Then, the combined transform resulting from motion correction, eddy-current correction and anatomical alignment was applied to the raw diffusion data. The table of gradient directions was appropriately adjusted to fit the resampled diffusion data (Leemans and Jones, 2009).

##### 2.3.2.2. Voxel level diffusivity modeling

Diffusion at the voxel level was modelled twice: First, we used the standard tensor model to calculate, within each voxel, the commonly used diffusion indices (e.g., FA and MD). These tensor-derived values were later entered into the statistical analyses along the tracts of interest (see section 2.4.). However, to overcome known limitations in the tensor model, in particular its inability to account for multiple orientations within the same voxel (Jones et al., 2013), our whole-brain tractography was based on CSD modelling. The CSD model resolves crossing fibers by estimating an orientation density function for each voxel, thereby improving the anatomical accuracy of tractography (Tournier et al., 2004, 2007).

###### Tensor model fitting

Tensors were fit to the diffusion measurements for each voxel in the aligned volume, using a robust least-squares algorithm, Robust Estimation of Tensors by Outlier Rejection (RESTORE), which removes outliers at the tensor estimation step (Chang et al., 2005). Tensor modelling was carried out using ‘mrDiffusion’ (https://github.com/vistalab/vistasoft). In each voxel, the estimated tensor eigenvalues (λ1, λ2, λ3) were used to calculate the diffusion metrics of interest (note that these tensors were *not* used by the probabilistic tractography algorithm). Specifically, Fractional anisotropy (FA) was calculated as the normalized standard deviation of the eigenvalues (Basser and Pierpaoli, 1996); Mean diffusivity (MD) was calculated as the average of all three eigenvalues; Axial diffusivity (AD) refers to the first eigenvalue (λ1), reflecting the diffusivity along the principal axis. Lastly, radial diffusivity (RD) was calculated as the average diffusivity along the second and third axes (λ2, λ3).

###### CSD model fitting

We used the mrTrix3 toolbox (https://www.mrtrix.org; Tournier et al., 2019) to calculate the CSD diffusion model. Diffusion response functions were estimated using the *dwi2response* command, implementing the *dhollander* algorithm (Dhollander et al., 2016; Dhollander and Connelly, 2016). This algorithm estimates separate response functions for white matter, gray matter and cerebrospinal fluid (CSF) based on single-shell diffusion-weighted data. The white matter and the CSF response functions were then used to estimate the fiber orientation distributions (FOD) based on constrained spherical deconvolution (CSD) with up to eight spherical harmonics (lmax = 8) (Tournier et al., 2004, 2007), using the *dwi2fod* command with the *msmt_csd* algorithm (Jeurissen et al., 2014). These FODs were then used for probabilistic tractography.

##### 2.3.2.3. Fiber tractography

Probabilistic whole brain tractography was carried out using the mrTrix3 command *tckgen* with the default iFOD2 tracking algorithm (improved 2^nd^-order integration over fiber orientation distributions; Tournier et al., 2010). Tracking was initialized from 500000 random seeds, placed within a whole brain white matter mask. The mask was generated from each subject’s structural T1 image using the *5ttgen* script, which performs whole brain segmentation utilizing FSL tools (Smith et al., 2004). The specific tracking parameters were as follows: FOD amplitude threshold 0.1, maximum angle between successive steps 45°, step size 0.85mm. Minimum and maximum streamline length values were set to 50mm and 200mm, respectively. Streamlines that extended beyond the white matter mask were truncated. The resulting whole brain tractogram was then used for automated tract segmentation.

For a full description of the deterministic tractography procedure see Supplementary material and our prior publication (Yablonski and Ben-Shachar, 2020).

##### 2.3.2.4. Tract identification and segmentation

We used automatic tract segmentation in order to identify each tract in the native space of each participant (see below for details). Based on our previous findings linking morphological processing and ventral-stream tracts (Yablonski et al., 2019) we targeted the IFOF, ILF, and the UF. Since any fluency task also involves speech production, we included additional dorsal tracts of interest: the fronto-temporal and fronto-parietal segments of the arcuate fasciculus (AF-ft and AF-fp, respectively), and the frontal aslant tract (FAT). Ventral and dorsal tracts were identified bilaterally in each individual’s native space.

Tract segmentation was carried out using the Automatic Fiber Quantification software package (AFQ; https://github.com/yeatmanlab/AFQ) (Yeatman et al., 2012b) implemented in Matlab 2012b. In accordance with this method, each tract is identified automatically by intersecting the whole brain fiber group with two waypoint ROIs, which are universally defined on a template, and then transformed to each individual’s native space. The ROIs were automatically transformed from the Montreal Neurological Institute (MNI) T2 template into each individual’s native space, using an estimated non-linear transformation. Individual waypoint ROIs were then intersected with the whole brain tractogram to isolate the tracts of interest. The resulting tracts were cleaned automatically, using a statistical outlier rejection algorithm that removed streamlines that were more than 3 standard deviations longer than the mean tract length, or that deviate by more than 4 standard deviations in distance from the tract core. This process was iterated 5 times for each tract. Note that these parameters are more restrictive than the default proposed in (Yeatman et al., 2012b) in order to reduce the amount of spurious streamlines that characterize probabilistic tractography.

Individual tracts were visually inspected using Quench, an interactive 3D visualization tool (Akers, 2006); http://web.stanford.edu/group/vista/cgi-bin/wiki/index.php/QUENCH). Visual inspection of the tracts revealed that the automatically cleaned AF-fp and UF contained stray streamlines that did not fit their anatomical definition. We thus employed stricter cleaning criteria to these tracts, as in our prior study (Yablonski and Ben-Shachar, 2020). Specifically, for the AF-fp, we excluded streamlines that were 1 standard deviation longer than the mean tract length, as well as streamlines that deviate by more than 4 standard deviations from the tract core. For the UF, we excluded any streamlines passing through an axial ROI at the level of the body of the corpus callosum (z = 22).

For each segmented tract, we delineated a diffusion profile by calculating the diffusion indices (FA, MD, AD, RD) at 100 equidistant nodes along the portion of the tract enclosed by the two ROIs. Within each tract, nodes are numbered from anterior (node 1) to posterior (node 100), except for the FAT where nodes are numbered from inferior (node 1) to superior (node 100). At each node, AFQ generates a weighted average of the diffusivity parameters in the streamlines belonging to the same tract. By restricting the analysis to the portion of the tract between the two waypoint ROIs, and by weighting more heavily streamlines that are closer to the core of the tract, our analyses focus on core white matter regions where tract trajectories are highly consistent across individuals, and more reliable within individual (Yeatman et al., 2012b). A different approach was applied to the UF, where diffusivity measures were analyzed along the entire trajectory, because the segment enclosed by the two UF ROIs misses the entire frontal portion of this tract. Statistical analyses were carried out on the resulting FA and MD profiles extracted from each tract as described below.

#### 2.3.3. CHARMED analysis

Preprocessing steps were carried out on the b=1000 s/mm^2^ shell data of the CHARMED sequence using ExploreDTI version 4.8.6 (Leemans et al., 2009). First, data were denoised using a regularization procedure and corrected for eddy currents and motion artifacts. The b-matrix (which includes the adjusted b-values in all gradient directions) was reoriented accordingly (Leemans and Jones, 2009). Then, a tensor model was fit for each voxel. The resulting FA map, diffusion tensors and the first eigenvectors were exported for the next processing steps, which were carried out in Matlab 2015b. The high b-value images were corrected for eddy currents and motion artifacts using the UNDISTORT method (Ben-Amitay et al., 2012). Finally, CHARMED analysis was performed using in-house software (Assaf and Basser, 2005; Tavor et al., 2013), resulting in individual maps of the volume fraction of the restricted compartment (FR). FR maps were aligned and registered to the anatomical T1 image and analyzed as an additional map within AFQ framework. FR maps of 3 subjects could not be properly aligned and were therefore not included in analyses of FR data.

### 2.4. Brain-behavior correlation analyses

For each of the tracts of interest, we extracted FA and MD values from each node along the tract profile. The Shapiro-Wilk test indicated that the distribution of FA and MD, averaged along each tract, deviated from normality. We therefore applied an outlier exclusion criterion that removed tracts whose average FA or MD deviated by more than 2.5 standard deviations from the group mean. We then calculated two-tailed Pearson’s correlation coefficients between FA and MD in each node of the tract profile and morpheme-based fluency scores. Significance was corrected for 100 comparisons (representing 100 nodes along the tract) using a nonparametric permutation method, yielding a family-wise error (FWE) corrected alpha value of 0.05 (Nichols and Holmes, 2003). Across the tracts, we controlled the false discovery rate (FDR) at a level of 5% (Benjamini and Hochberg, 1995). We then followed up on significant correlations by calculating multiple regression models controlling for standard measured of verbal fluency. Specifically, within each significant cluster, we calculated multiple linear regression models predicting FA (or MD), based on morpheme-based fluency, category-based fluency and letter-based fluency, as well as age. Regression analyses were carried out in R and visualized using the *ggplot2* package (R Core Team, 2019; Wickman, 2016).

To interpret the direction of the observed correlations post-hoc, we extracted AD and RD values from clusters where FA or MD were significantly correlated with morpheme-based fluency. We then calculated two-tailed Pearson’s correlations between morpheme-based fluency and mean AD or mean RD in the same clusters. Throughout our analyses we used two-tailed statistics, because previous literature reports both positive and negative correlations between FA and reading related skills (Frye et al., 2011; Welcome and Joanisse, 2014; Yeatman et al., 2012a).

To further our insight into the biological factors underlying the effects, we extracted FR values from clusters where FA or MD effects were found. Unlike AD and RD, FR is an independent measure calculated over data obtained in a separate scan (see above). We calculated two-tailed Pearson’s correlations between morpheme-based fluency and mean FR extracted from significant clusters. Lastly, to examine whether correlations with FR along the tract profiles reveal a similar picture to the one obtained using tensor-derived metrics (FA, MD), we calculated two-tailed Pearson’s correlation coefficients between morpheme-based fluency scores and FR in each node along each of the tracts. We accounted for multiple comparisons in a similar manner as we did for FA and MD: a nonparametric family-wise error correction across 100 nodes in each tract, then controlling the FDR across the tracts at a level of 5%.

## 3. Results

### 3.1. Verbal fluency scores

Individual performance in the fluency tasks is reported in Figure 1 and Supplementary Table S1. Two subjects were excluded as outliers (see *Methods* for exclusion criterion): One subject was excluded from the root-based, pattern-based and category-based fluency tasks, and another subject was excluded from the letter-based fluency task. The Shapiro-Wilk test confirmed that the remaining scores (N=44 for each task) were normally distributed. Consistent with previous literature (Blecher et al., 2019; Kavé, 2005; Shao et al., 2014), letter-based fluency scores were lower than category-based fluency scores, and the two tasks were correlated (r = 0.53, p < 0.05, after controlling the FDR across all behavioral measures at q< 0.05; See Figure 1). Letter-based fluency was also strongly correlated with pattern-based fluency (r = 0.6, p < 0.05). Interestingly, performance on the timed single word reading test (Shatil, 1997) was positively correlated with all fluency tasks, except for root-based fluency, suggesting that root-based fluency requires some distinct processes. The full inter-correlation matrix is reported in Supplementary Table S3.

### 3.2. Brain-behavior correlations

The twelve tracts of interest were successfully detected by probabilistic tractography in all 45 participants. Figure 2 depicts the identified tracts in three representative subjects. The Shapiro-Wilk test confirmed that the mean tract FA and MD in each of the tested tracts were normally distributed following outlier exclusion (see *Methods*). In total, 0-2 subjects were excluded as outliers from subsequent analyses of each tract (actual sample sizes included in neurocognitive correlations for each tract are reported in Tables S4 and Table S5 for FA and MD, respectively). The deterministic tractography pipeline failed to detect the right AF-ft, left FAT and right FAT in several participants (see Supplementary Tables S4-S5), as previously documented (Catani et al., 2007; Vanderauwera et al., 2017; Yablonski et al., 2019; Yeatman et al., 2011). Direct comparisons between the diffusion metrics obtained from each tract using the two analysis procedures are reported in Supplementary Figures S1-S3.

**Figure 2.**
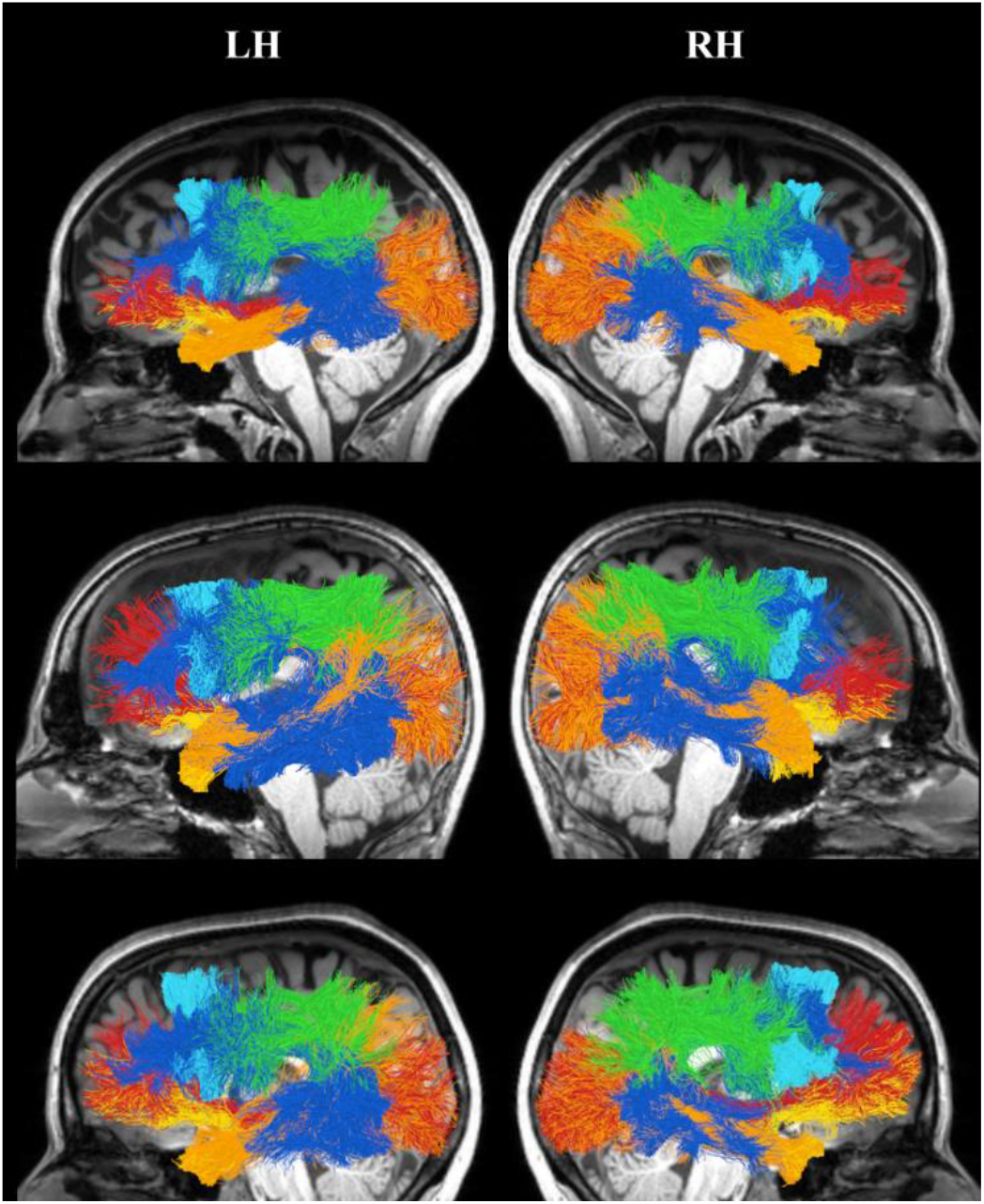
Dorsal and ventral tracts of interest. Shown are the bilateral tracts identified in three representative subjects (female, 22y; male, 30y; female, 24y), overlaid on their midsagittal T1 image. The ventral tracts identified are the inferior longitudinal fasciculus (orange), the inferior fronto-occipital fasciculus (red) and the uncinate fasciculus (yellow). The dorsal tracts identified are the fronto-temporal (blue) and fronto-parietal (green) segments of the arcuate fasciculus, and the frontal aslant tracts (cyan). LH-left hemisphere, RH-right hemisphere.

#### 3.2.1. Root-based fluency is associated with diffusivity in ventral tracts

Based on our previous findings (Yablonski et al., 2019; Yablonski and Ben-Shachar, 2020), we hypothesized that morpheme-based fluency is associated with ventral language tracts, bilaterally. This prediction was born out. Indeed, an analysis of the correlations along the tracts revealed that root-based fluency was negatively correlated with MD in the IFOF and ILF, bilaterally (see Figure 3 and Table 1; all significant findings reported here are significant at p < 0.05, FWE corrected for 100 comparisons along each tract, controlling the FDR at q < 0.05 across the ventral and dorsal tracts of interest). The locations of the significant clusters along the ventral tracts are visualized in Figure 3 (panels a, c, e, g). The distributions of individual values and the patterns of covariation between root-based fluency and MD within these significant clusters are also visualized in Figure 3 (panels b, d, f, h). In addition, root-based fluency was positively correlated with FA in a cluster in the right ILF (Figure 4; panels a, b). Similar MD clusters were found when the ventral tracts were reconstructed using tensor modeling and deterministic tractography (see Supplementary Figure S4 and Table S6). No significant correlations were found between pattern-based fluency and diffusion properties in any of the ventral tracts.

**Figure 3.**
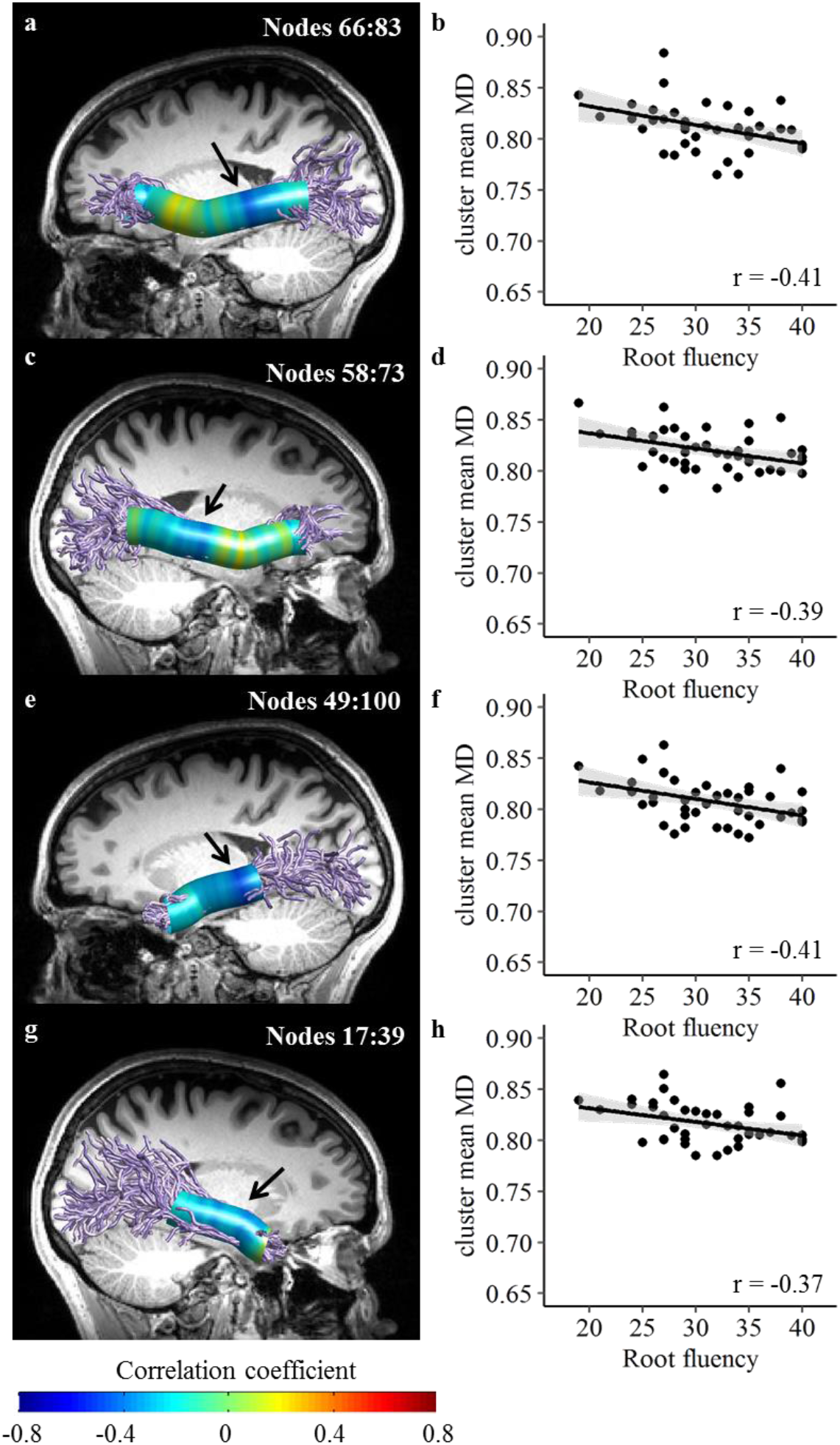
Root based fluency is negatively correlated with MD in bilateral ventral tracts. (a, c, e, g) Pearson’s correlation coefficients are visualized in 100 nodes along the left IFOF (a), right IFOF (c), left ILF (e) and right ILF (g). Black arrows denote the location of significant clusters after family-wise error correction across the 100 nodes. (b, d, f, h) Scatter plots showing the association between root-based fluency (number of words) and the mean MD in the significant cluster of nodes, in left IFOF (b), right IFOF (d), left ILF (f) and right ILF (h). Black lines represent the best linear fit, surrounded by the 95% confidence interval (shaded area). These scatter plots are shown for visualization purposes, significance is calculated along the trajectory of the tracts. IFOF-inferior fronto-occipital fasciculus. ILF-inferior longitudinal fasciculus.

**Table 1.** Root-based fluency is correlated with diffusion properties in ventral and dorsal tracts, bilaterally. Reported are clusters of nodes showing significant Pearson’s correlations with Root-based fluency, family-wise error corrected for 100 nodes. Significant clusters were followed up with multiple regression models predicting cluster mean FA or MD from root-based fluency, as well as category-based fluency, letter-based fluency and age. We report the R squared of each regression model and the significance level of the root fluency predictor. The contribution of the additional predictor variables was non-significant (p > 0.05) in all the models tested. *p < 0.05, FDR corrected for 8 clusters. § p < 0.05 for the root-based fluency predictor within each regression model

|  | cluster<br>location<br>(Nodes) | r | p | CI (95%) | Regression<br>Full model<br>R<sup>2</sup> | Root<br>fluency<br>p |
| --- | --- | --- | --- | --- | --- | --- |
| **FA** |  |  |  |  |  |  |
| Right ILF | 33 – 53 | 0.41 | 0.006* | [0.11, 0.63] | 25% | 0.002§ |
| Left FAT | 29 – 45 | 0.46 | 0.002* | [0.21, 0.65] | 23% | 0.004§ |
| **MD** |  |  |  |  |  |  |
| Left IFOF | 66 – 83 | -0.41 | 0.007* | [-0.56, -0.17] | 26% | 0.004§ |
| Right IFOF | 58 – 73 | -0.39 | 0.011* | [-0.63, -0.07] | 22% | 0.010§ |
| Left ILF | 49 – 100 | -0.41 | 0.007* | [-0.60, -0.13] | 20% | 0.008§ |
| Right ILF | 17 – 39 | -0.37 | 0.015* | [-0.57, -0.02] | 31% | 0.005§ |
| Left FAT | 26 – 63 | -0.40 | 0.007* | [-0.65, -0.05] | 23% | 0.015§ |
| Right FAT | 15 – 59 | -0.50 | 0.001* | [-0.71, -0.19] | 27% | 0.005§ |

**Figure 4.**
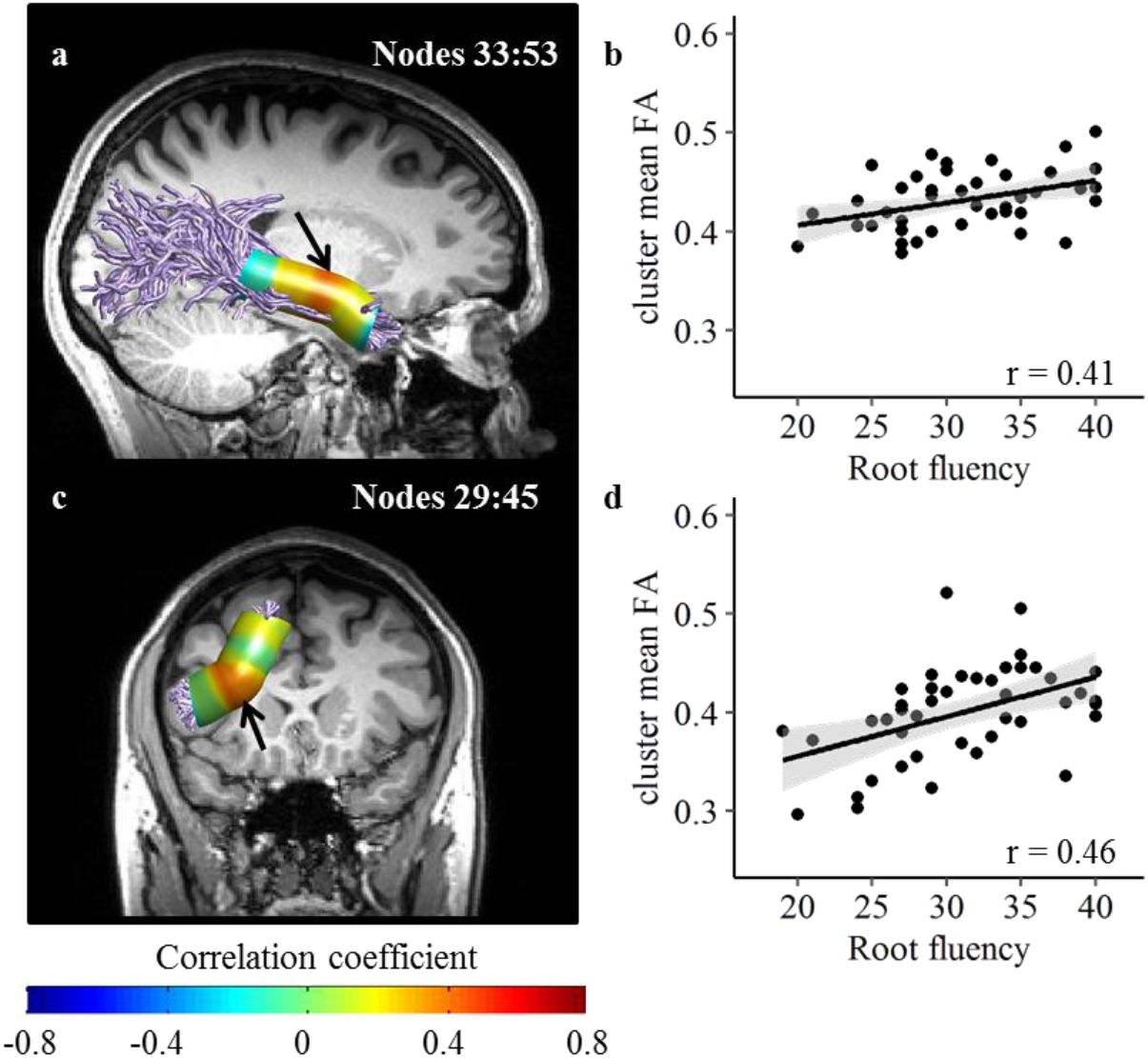
Root-based fluency is positively correlated with FA in the right ILF and left FAT. (a, c,) Pearson’s correlation coefficients are visualized in 100 nodes along the right ILF (a) and left FAT (c). Black arrows denote the location of significant clusters after family-wise error correction across the 100 nodes. (b, d) Scatter plots showing the association between root-based fluency (number of words) and the mean FA in the significant cluster of nodes, in the right ILF (b) and left FAT (d). Black lines represent the best linear fit, surrounded by the 95% confidence interval (shaded area). These scatter plots are shown for visualization purposes, significance is calculated along the trajectory of the tracts. ILF-inferior longitudinal fasciculus. FAT-frontal aslant tract.

#### 3.2.2. Root-based fluency is further associated with diffusivity in dorsal tracts

Since verbal fluency tasks involve speech production, we expected to find associations in dorsal tracts as well. Indeed, root-based fluency was negatively correlated with local MD values in the bilateral FAT (Figure 5; Table 1). In the left FAT, root-based fluency was also positively correlated with local FA values, in a cluster of nodes overlapping the cluster that showed MD effects (Figure 4; panels c, d). No significant correlations were found between pattern-based fluency and diffusion properties in any of the dorsal tracts. Interestingly, when dorsal tracts were reconstructed using deterministic tractography, no correlations were found in the left or right FAT, but instead, significant correlations were found in the left AF-ft. This correlation was weaker than the correlations found in the ventral tracts (see Supplementary Figure S5 and Table S6). The correlations found in the dorsal tracts are therefore more sensitive than the ventral tracts to the specific choice of analysis approach.

**Figure 5.**
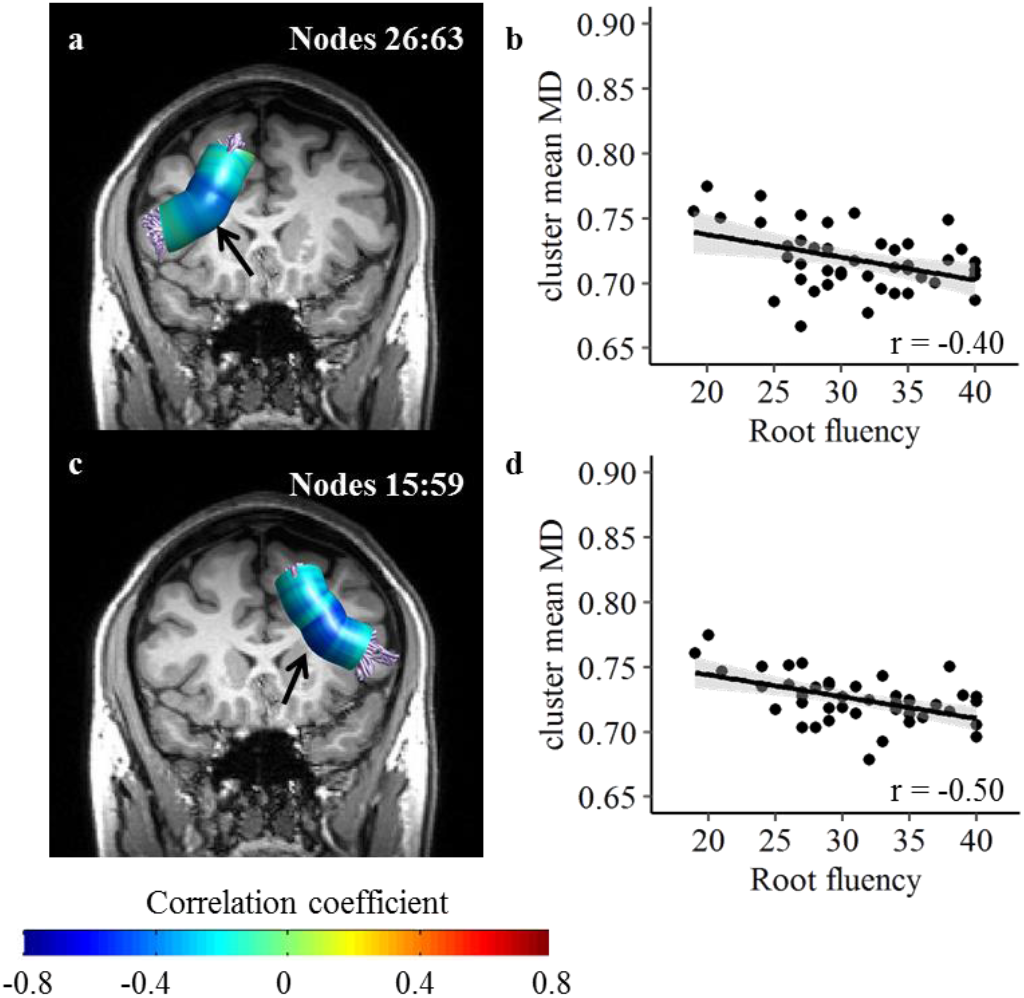
Root-based fluency is negatively correlated with MD in the frontal aslant tract, bilaterally. (a, c,) Pearson’s correlation coefficients are visualized in 100 nodes along the left (a) and right FAT (c). Black arrows denote the location of significant clusters after family-wise error correction across the 100 nodes. (b, d) Scatter plots showing the association between root-based fluency (number of words) and the mean MD in the significant cluster of nodes, in the left (b) and right FAT (d). Black lines represent the best linear fit, surrounded by the 95% confidence interval (shaded area). These scatter plots are shown for visualization purposes, significance is calculated along the trajectory of the tracts. FAT-frontal aslant tract.

For completeness, we also calculated the correlations between root-based fluency and FA or MD averaged across the length of each tract, but none of these reached significance in the probabilistic tractography approach. However, using deterministic tractography with tensor modeling, the same analysis produced significant correlations between root-based fluency and MD in the bilateral IFOF and bilateral ILF (see Supplementary Table S5), thus providing additional support to the main finding, that generalizes our previously reported morphological associations in the bilateral ventral language tracts, across modality and task.

#### 3.2.3. Specificity of the associations with root-based fluency

To examine the specificity of the associations reported so far, we followed up on significant clusters with multiple linear regression analyses. For each cluster that showed a significant correlation with root-based fluency, we calculated a regression model predicting cluster mean MD (or FA) based on root-based fluency, letter-based fluency, category-based fluency, and age. In all of the clusters reported above, root-based fluency was a significant predictor of MD or FA (p < 0.05, see Table 1), while the contribution of the other predictors was non-significant (p > 0.1; except for the right ILF where the contribution of letter-based fluency was near significance, p = 0.0504). In other words, the correlations with root-based fluency remained significant when age and standard fluency measures were accounted for. These models explained between 20-31% of the variance in cluster mean MD or FA. These results are reported in Table 1 and visualized as scatterplots in Supplementary Figures S6-S7. Note that the scatterplots in Figures S6 parallel those in Figure 3 and Figure 5, but show the regression residuals instead of the raw MD and fluency values. We chose to visualize raw values in the main figures (rather than residuals) because the raw values have units and are easier to interpret.

#### 3.2.4. Follow-up analyses with diffusivity subcomponents

To elucidate the source of the effects found with FA and MD, we extracted AD and RD values in the clusters that showed significant correlations with root-based fluency. We then examined, separately, the pattern of association between each of these measures and root-based fluency. As shown in Table 2, no correlations were found with AD. However, root-based fluency was negatively correlated with RD in the left IFOF, right ILF, and left and right FAT (p < 0.05, controlling the FDR across the 8 significant clusters). We next examined the association between root-based fluency and FR values, which are derived from a separate scan, and provide an additional independent index of estimated fiber density. We found that root-based fluency was positively correlated with FR in the left IFOF (Figure 6), but not in any of the other significant clusters. This finding suggests that the underlying factor driving the correlation in the left IFOF is different than in the other tracts. This pattern was consistent when RD and FR were extracted from tracts reconstructed using deterministic tractography (Supplementary Figure S8 and Table S7).

**Table 2.**
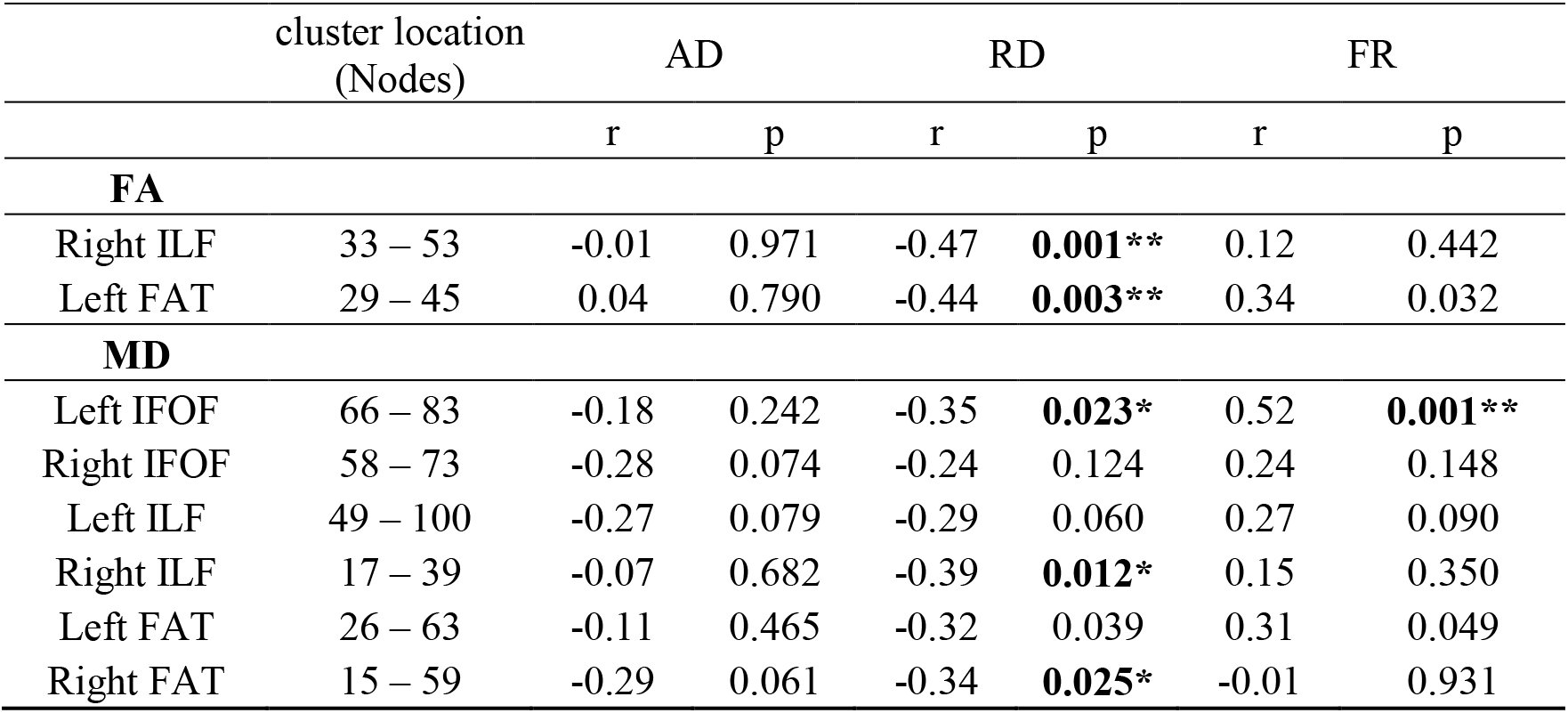
Root-based fluency is correlated with FR in the left IFOF. For each significant cluster we extracted the mean value of axial diffusivity (AD), radial diffusivity (RD) and the restricted diffusion fraction from the CHARMED model (FR), an index of fiber density. Although clusters in several tracts show a significant correlation with RD, correlations with FR are found in the left IFOF alone. Reported are Pearson’s correlation coefficients between each diffusion component and root fluency. *p < 0.05, FDR corrected for 8 clusters. ** p < 0.05, FDR corrected for 24 comparisons (8 clusters * 3 measurements)

**Figure 6.**
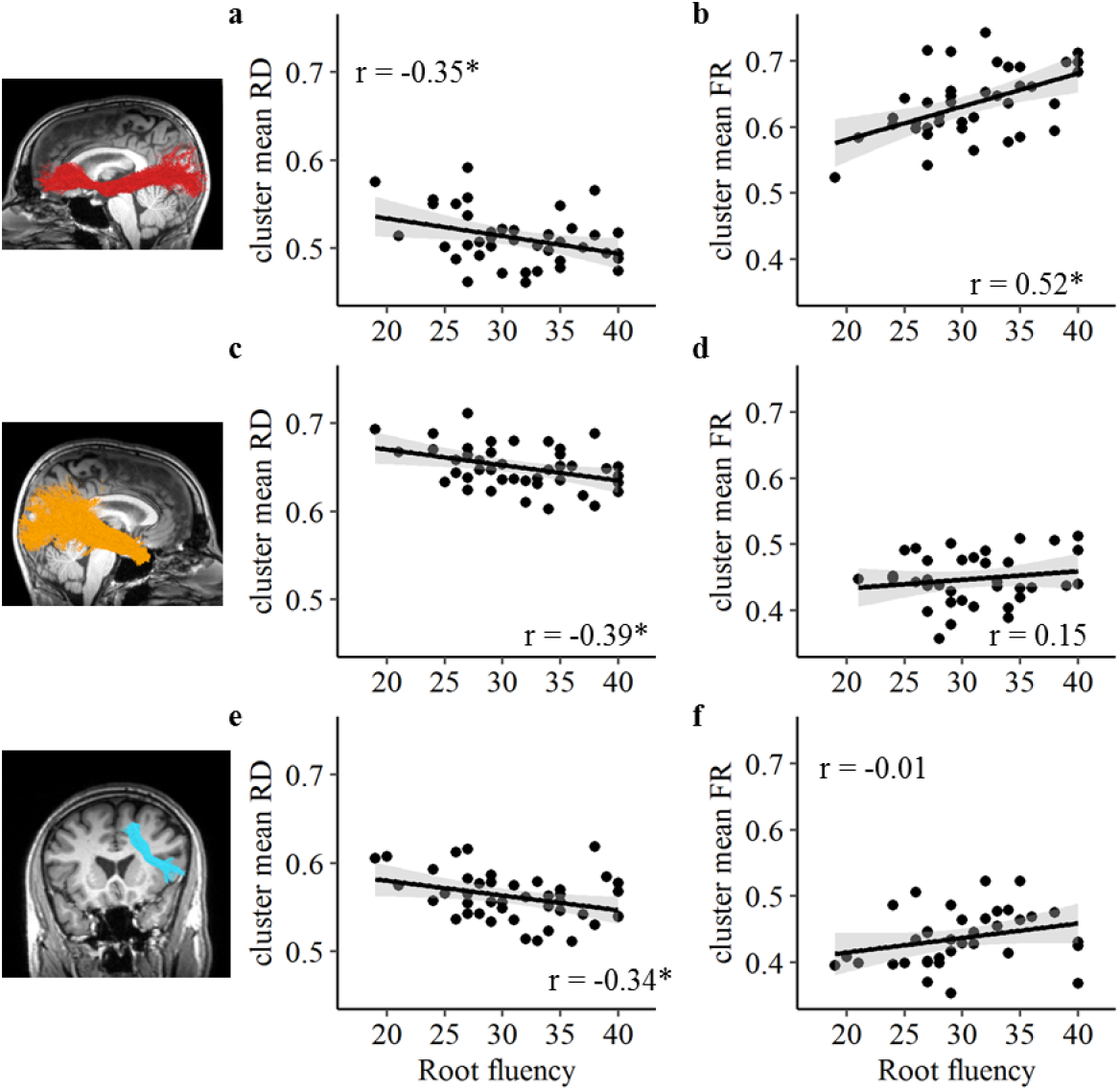
Root-based fluency is correlated with RD in several tracts, yet correlations with FR are found only in the left IFOF. Scatter plots show the association between root-based fluency and mean radial diffusivity (RD; panels a, c, e) or mean diffusivity in the restricted compartment (FR; panels b, d, f) within clusters showing a significant correlation between RD and root-based fluency (see Table 2 for a full report of the correlations with RD and FR in all the significant clusters). Tractograms on the left demonstrate the relevant pathway in a single participant. *p < 0.05, FDR corrected for the 8 clusters that showed significant correlations with root-based fluency.

#### 3.2.5. FR analysis along the tract uncovers additional correlations

To examine the utility of including FR as an additional structural index, we next computed the correlations between root-based fluency and local FR values along each tract of interest. This analysis revealed significant positive correlations in the left IFOF and left ILF (Figure 7 and Table 3; see Supplementary Table S8 for FR values averaged along the tracts). In both tracts, the significant cluster overlaps the location of the clusters found with MD. Surprisingly, this analysis also uncovered a positive correlation between root-based fluency and FR in a cluster in the left AF-ft, a dorsal tract (Figure 8; Table 3). This finding suggests that FR may provide increased sensitivity for detecting neurocognitive correlations, beyond those observed with tensor derived measures (e.g., FA, MD). Multiple regression models verified that root-based fluency was a significant predictor of FR, over and above the contribution of letter-based fluency, category-based fluency and age (p < 0.05, FDR corrected; Table 3). The contribution of the additional predictors was non-significant in all the regression models tested (p > 0.05), except for the left AF-ft, where letter-based fluency added a significant contribution (p = 0.013), in alignment with this tract’s involvement in phonological operations (e.g., Yeatman et al., 2011). These findings were replicated using the deterministic tractography pipeline (Supplementary Table S9; Figures S9-S10).

**Figure 7.**
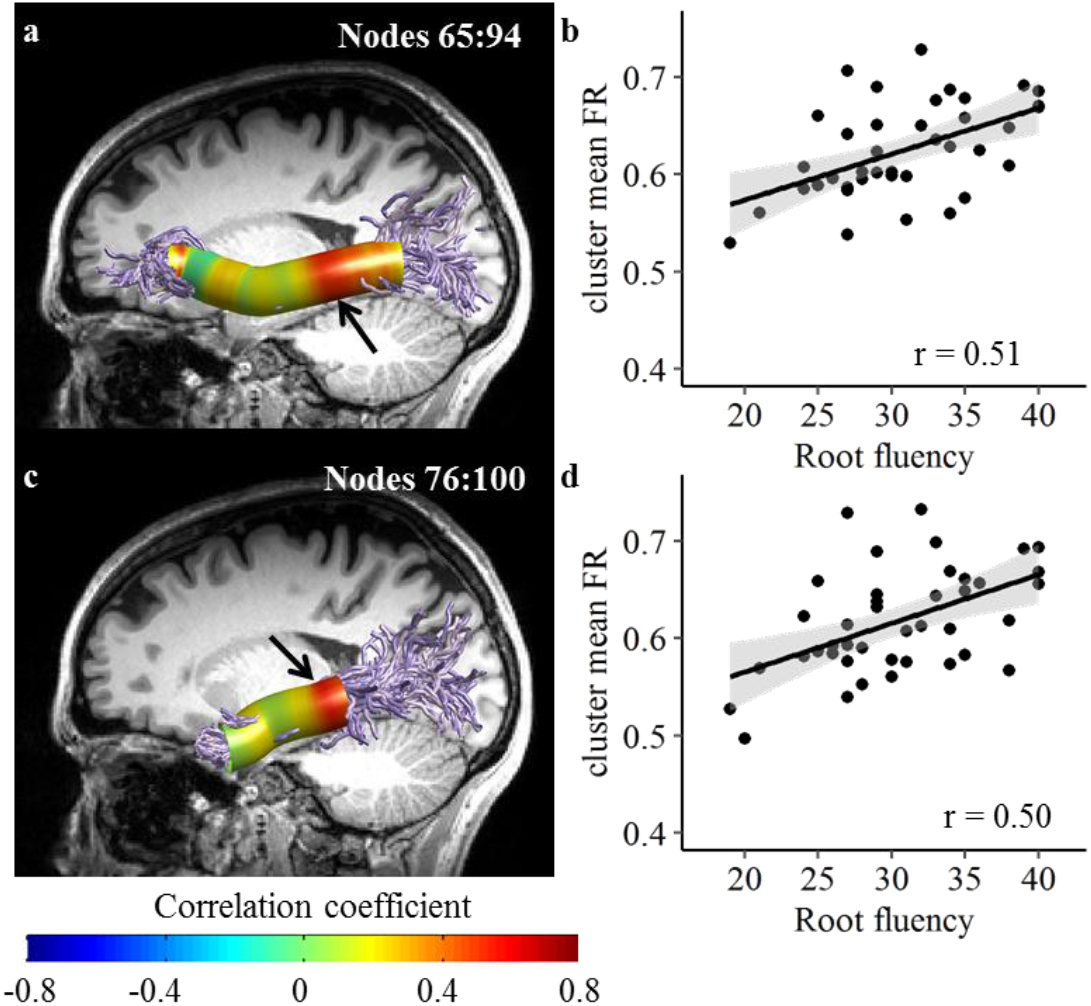
Analysis of FR along the tracts reveals selective correlations with ventral tracts in the left hemisphere. Root based fluency is positively correlated with FR in significant clusters in the left IFOF (a-b) and left ILF (c-d). No significant clusters were found in the right hemisphere tracts that showed significant FA or MD effects. Black lines represent the best linear fit, surrounded by the 95% confidence interval (shaded area). These scatter plots are shown for visualization purposes, significance is calculated along the trajectory of the tracts. IFOF-inferior fronto-occipital fasciculus. ILF-inferior longitudinal fasciculus.

**Table 3.** Diffusivity in the restricted compartment (FR) is correlated with root-based fluency and pattern-based fluency. Reported are clusters of nodes showing significant Pearson’s correlations between FR and Root-based fluency or Pattern-based fluency, family-wise error corrected for 100 nodes. Significant clusters were followed up with multiple regression models predicting cluster mean FR from root-based or pattern-based fluency, as well as category-based fluency, letter-based fluency and age. We report the R squared of each regression model and the significance level of the predictor of interest, root or pattern. The contribution of the additional predictor variables was non-significant in all the models tested (p > 0.05), except for the left AF-ft, where letter-based fluency made a significant contribution (p = 0.013). *p < 0.05, FDR corrected for 4 clusters. §p < 0.05 for the morpheme-based fluency predictor (root or pattern as indicated) within each regression model

|  | cluster location<br>(Nodes) | r | p | CI (95%) | Regression |  |
| --- | --- | --- | --- | --- | --- | --- |
|  |  |  |  |  | Full model<br>R<sup>2</sup> | Root fluency<br>p |
| Root-based fluency |  |  |  |  |  |  |
| Left IFOF | 65 – 94 | 0.51 | 0.001* | [0.23, 0.72] | 40% | 1.4x10<sup>-4</sup>§ |
| Left ILF | 76 – 100 | 0.50 | 0.001* | [0.22, 0.72] | 32% | 2.5x10<sup>-4</sup>§ |
| Left AF-ft | 81 – 100 | 0.62 | 1.5x10<sup>-5</sup>* | [0.41, 0.78] | 51% | 1.5x10<sup>-5</sup>§ |
| Pattern-based fluency |  |  |  |  |  |  |
| Right UF | 67 – 100 | 0.42 | 0.007* | [0.20, 0.61] | 25% | 0.003§ |

**Figure 8.**
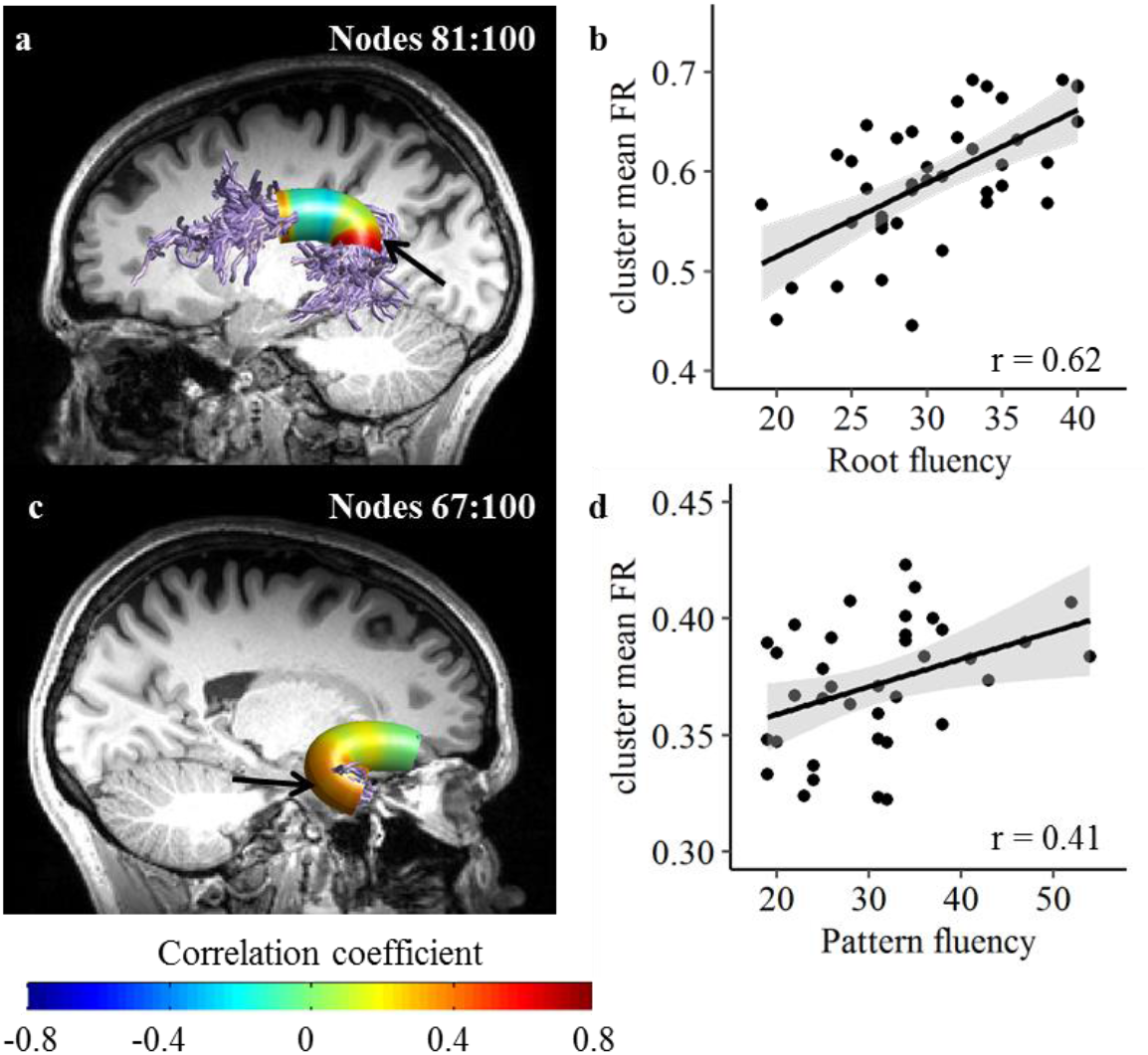
Analysis of FR along the tracts uncovers correlations that were not detected using traditional diffusion metrics. Root based fluency is correlated with FR in a significant cluster in the left AF-ft (a-b). Pattern based fluency is correlated with FR in a significant cluster in the right UF (c-d). Black lines represent the best linear fit, surrounded by the 95% confidence interval (shaded area). These scatter plots are shown for visualization purposes, significance is calculated along the trajectory of the tracts. AF-ft-fronto-temporal segment of the arcuate fasciculus. UF-uncinate fasciculus.

After observing that FR may detect correlations not discovered otherwise, we looked into correlations between FR and pattern-based fluency, which was not correlated with FA or MD in any of the tracts of interest. We found a significant cluster of positive correlation between pattern-based fluency and FR in the right UF (Figure 8). A multiple regression model confirmed that pattern-based fluency was the only significant predictor of FR values in this cluster (Table 3). Interestingly, when we investigated correlations with FR values extracted from deterministic tractography tracts, pattern-based fluency was correlated with FR in the right AF-ft (Supplementary Table S9; Figure S10).

Lastly, to understand the relationship between FR and the traditional diffusion metrics, we conducted an exploratory analysis of the correlations between FR and FA, MD and RD, averaged across the tracts that showed significant effects. This analysis revealed that FR is positively correlated with FA, in line with previous reports (De Santis et al., 2014). This was mirrored by negative correlations between FR and MD, and between FR and RD (Supplementary Figures S11-S13). Interestingly, the strength of the correlations varied greatly across the different tracts, indicating weaker coupling between FR and tensor-derived metrics in the AF-ft and FAT compared with the IFOF and ILF. Indeed, AF-ft is where we document the most striking divergence between standard diffusivity measures (MD in particular) and FR.

## 4. Discussion

The goal of the current study was to examine whether the involvement of ventral white matter pathways in morphological processing generalizes across modalities. We found that performance in an oral morpheme-based word production task was correlated with properties of the bilateral ventral tracts associated with language and reading. This finding complements previous findings relating these tracts to morphological processing applied to visually presented written stimuli (Yablonski et al., 2019; Yablonski and Ben-Shachar, 2020). In addition, morpheme-based fluency was correlated with the bilateral FAT, a dorsal tract previously associated with speech production (Catani et al., 2013). Importantly, follow-up analyses confirmed that the correlations observed with morpheme-based fluency remained significant when controlling for performance in standard fluency tasks. Lastly, we found that some of the correlations may be explained by FR, a novel index of estimated fiber density, and demonstrated that incorporating FR as an additional variable can uncover effects beyond those obtained using tensor-derived metrics.

The current study makes two major contributions: First, we show that an oral task that requires morphological processing recruits both ventral pathways and dorsal pathways, bilaterally. We interpret the correlations between morpheme-based fluency and the ventral tracts as further confirmation for these tracts’ involvement in morphological processing, generalizing previous findings across task, input modality and response modality. Further, the correlations found in the ventral tracts were robust to the choice of analysis approach (compare, for example, Figure 3 with supplementary Figure S4). In contrast, the correlations between morpheme-based fluency and the microstructure of the FAT may be more specific to the production task used in the current study. This demonstrates that neurocognitive correlations are modulated not only by the cognitive processes in question (i.e., morphological processing), but also by the tasks used to probe them. The effects in the FAT were more susceptible to analysis choices and only emerged when probabilistic tractography was employed. Together, these findings highlight the influence of task selection and experimental design on the obtained effects, in addition to the cognitive construct in question.

Second, the current study makes an important contribution to the literature by incorporating FR, a novel diffusion-based index of white matter microstructure. To date, only a handful of studies have investigated correlations between FR and cognitive skills (Metzler-Baddeley et al., 2017; Tavor et al., 2013). By analyzing FR in addition to standard diffusion metrics, we were able to dissociate the effect in the left IFOF from the rest of the tracts that demonstrated correlations with morpheme-based fluency. This implies that the underlying tissue property driving the effect in the left IFOF may be different than in the other tracts. While FA and MD are influenced by a multitude of tissue properties, FR is more selective to the intra-axonal diffusion component and is generally considered an index of fiber density (De Santis et al., 2014). Furthermore, using FR we uncovered additional correlations that could not be detected using FA or MD, specifically in the left AF-ft and right UF. This is in line with previous findings showing that FR is more sensitive than FA or MD (De Santis et al., 2017; Tavor et al., 2013). Our exploratory analyses further revealed that the coupling between FR and FA is modulated by structural organization, corroborating previous observations (De Santis et al., 2014). For example, FA and FR are more tightly linked in tracts that are directionally cohesive, like the IFOF, compared with tracts that include multiple fiber orientations in a complicated architecture, like the AF-ft (Supplementary Figures S11-S13). This suggests that FR is less susceptible than FA or MD to the effects of directional coherence, and taps more specifically onto local cellular properties. Our findings provide novel evidence for the potential of FR as a sensitive index for elucidating correlations between cognitive abilities and white matter microstructure.

### 4.1. Morpheme-based fluency is associated with ventral and dorsal tracts

In the current study, participants used auditorily presented morphological information as a cue for lexical search and oral word production. Although this task did not explicitly involve reading, performance was correlated with the bilateral IFOF and ILF, suggesting that their role in morpheme-mediated lexical access goes beyond their well-established involvement in orthographic processing (Horowitz-Kraus et al., 2014; Ozernov-Palchik et al., 2019; Vandermosten et al., 2012; Welcome and Joanisse, 2014; Yeatman et al., 2012a). Importantly, the associations between morpheme-based fluency and the ventral tracts remained significant when performance in standard verbal fluency tasks was accounted for, suggesting that the observed effects are somewhat specific to the morphological component of the task. These findings thus provide an important generalization across task, presentation modality and response modality, suggesting that ventral tracts are involved in amodal morpheme-mediated lexical access. This lends further support for the central role of morphological information in mapping form to meaning (Rastle, 2019), a process which has been associated with the ventral pathways (Friederici, 2012; Hickok and Poeppel, 2007; Saur et al., 2008).

Interestingly, morpheme-based fluency was also correlated with properties of the bilateral FAT, a dorsal tract connecting the inferior frontal gyrus (IFG) with the supplementary motor area (SMA) and pre-SMA (Catani et al., 2013; Thiebaut de Schotten et al., 2012). Several studies have found that the left FAT is critically involved in speech fluency in clinical populations (Blecher et al., 2019; Catani et al., 2013; Kronfeld-Duenias et al., 2016; Li et al., 2017; Mandelli et al., 2014). To the best of our knowledge, this is the first demonstration of this tract’s involvement in verbal fluency in healthy adults. Our regression analyses confirmed that the correlations between FAT microstructure and morpheme-based fluency were not explained by standard verbal fluency tests. The apparent specificity of the correlations with the morpheme-based task are in line with several case studies where surgical resection damaging the left FAT induced speech disruptions that were sensitive to grammatical constraints (Chernoff et al., 2019, 2018; Sierpowska et al., 2015). Of particular relevance is the case reported by Sierpowska and colleagues (2015), where damage to the left FAT elicited production errors characterized by inappropriate application of morphological rules. Taken together, these prior findings and the current results suggest that the role of the FAT in speech production may extend to high level language functions beyond its contribution to articulation and motor planning (Dick et al., 2019).

In the current study, morpheme-based fluency was associated both with the left and the right FAT. Although the right FAT has been relatively understudied compared with its left homolog, there is some evidence that the right FAT plays a role in developmental stuttering (Kronfeld-Duenias et al., 2016; Neef et al., 2018), suggesting an involvement in speech production. In contrast, no associations were found between the right FAT and verbal fluency in chronic stroke patients (Li et al., 2017). Considering these mixed findings and its cortical terminations, it has recently been postulated that the right FAT supports general inhibitory control (Dick et al., 2019). According to this view, the FAT supports the process of selecting a specific action program among several competitors that require the same motor resources. The left FAT does so specifically for speech, selecting one articulatory program and inhibiting others, while the right FAT is less specific to language and exerts inhibition in general motor planning. This intriguing possibility is partly supported by our current results, as morpheme-based fluency obviously requires participants to inhibit semantically and phonologically related words, as well as inflections of previous responses. However, we observed a similar pattern of results in the left and right FAT, therefore our current data do not support different functions for the left and right FAT in the context of verbal fluency.

Lastly, our secondary analyses suggested that another dorsal tract, the left AF-ft, is involved in morpheme-based fluency. Although morpheme-based fluency was not associated with the left AF-ft in the main analysis, a significant correlation emerged when this tract was reconstructed using the tensor-deterministic tractography procedure. Further, analysis of FR values along the AF-ft uncovered an additional cluster of correlation with morpheme-based fluency in this tract. Critically, multiple regression analyses revealed that letter-based fluency was an additional significant predictor in this cluster, in line with the well-established involvement of the left AF-ft in phonological processing (Saygin et al., 2013; Vanderauwera et al., 2015; Vandermosten et al., 2012; Yeatman et al., 2011). The correlation in the left AF-ft may reflect the auditory-motor mapping required in the morpheme-based fluency task, a process attributed to the dorsal language pathway (Hickok and Poeppel, 2007; Saur et al., 2008). Alternatively, the correlation in the left AF-ft may reflect its contribution to morphological processing per se. This interpretation is in accordance with the view of the left AF-ft as a central pathway for high-level linguistic processes, particularly supporting syntactic operations (Friederici, 2012; Skeide et al., 2016; Wilson et al., 2011). We consider the interpretation of the current findings in terms of task characteristics more likely, specifically because we did not find an association with morphological processing in the left AF-ft in response to written stimuli (Yablonski et al., 2019; Yablonski and Ben-Shachar, 2020). It should be noted that the finding in the left AF-ft was inconsistent across the different analytic procedures and structural indices, and should thus be considered with caution pending future replication.

To the best of our knowledge, this is the first study to look into the white matter correlates of morpheme-based fluency. Recently, a different task tapping into morpheme-based word production was selectively associated with the left ILF in children (Su et al., 2018). Interestingly, their task required participants to generate only two responses per target, and was thus less demanding in terms of working memory, inhibition and lexical search. This could explain the more restrictive pattern of results. Still, this finding provides supporting evidence for the involvement of the ventral tracts in morphological processing sampled with production tasks. Other studies reported associations between letter- or category-based fluency and properties of several white matter tracts, in clinical patients suffering from multiple sclerosis, aphasia or stroke (Blecher et al., 2019; Griffis et al., 2017; Keser et al., 2018; Li et al., 2017; but see Vallesi and Babcock (2020) for null results in healthy adults). Morpheme-based fluency may be more challenging than standard fluency tasks, as it requires using metalinguistic knowledge for lexical search and applying more restrictive response selection criteria. It is thus possible that morpheme-based fluency is better suited to study neurocognitive correlations in typical adults. Clearly, more research is needed to elucidate the white matter correlates of verbal fluency in the typical population.

### 4.2. Cognitive processes underlying the morpheme-based fluency task

Morpheme-based fluency requires participants to search the mental lexicon and orally produce words that share a morpheme with the target. According to models of speech production, the process of generating any spoken word consists of several steps: conceptual preparation, lexical selection of the appropriate word, morphological and phonological encoding, phonetic encoding, and finally articulation (Levelt et al., 1999). Notably, the model also highlights the importance of self-monitoring throughout the process. Although originally developed to account for performance in picture naming tasks, the same model can be applied to verbal fluency tasks, presuming that word generation in this case begins with lexical search according to the provided cue (Friedman et al., 1998; Indefrey and Levelt, 2000). In the morpheme-based fluency task, lexical search is guided by morphemes, which play an important role in the organization of the lexicon by forming a bridge between the semantic system and phonological word representations (Levelt et al., 1999). Morphemes are particularly central in the Hebrew language, where it was suggested that the organization of the mental lexicon is primarily governed by morphological principles (Deutsch et al., 1998; Frost et al., 2005, 2000). Recent findings indicate that morphemes guide lexical access in Hebrew word production, in addition to their well-established role in word perception (Deutsch, 2016; Deutsch and Malinovitch, 2016; Deutsch and Meir, 2011). Our findings provide further support for the importance of morphological units in lexical access across perception and production alike (Yablonski and Ben-Shachar, 2020, and current results).

Interestingly, verbal fluency tasks are classically administered as part of batteries evaluating executive functions (e.g., Kramer et al., 2014). Successful performance in verbal fluency requires, in addition to lexical search, response selection and inhibition of irrelevant responses, sustained attention and self-verification (Friedman et al., 1998). Indeed, multiple studies have found that verbal fluency tasks are correlated with executive functions, particularly with aspects of updating and working memory (Gustavson et al., 2019; Kavé and Sapir-Yogev, 2020; Shao et al., 2014; Unsworth et al., 2011; but see Whiteside et al., 2016). It was suggested that the cue used to prompt lexical search (i.e., letter or category) dictates different search strategies and exerts different levels of inhibition when generating responses. For example, category-based fluency supposedly capitalizes on the natural semantic organization of the lexicon, while letter-based fluency requires using less practiced strategies, as well as inhibiting semantically related words, making this task altogether more taxing (Basso et al., 1997). This notion is supported by consistent reports of better performance in category-based fluency compared with letter-based fluency (replicated here, and see Blecher et al., 2019; Kavé, 2005; Kavé and Knafo-Noam, 2015; Meinzer et al., 2009; Shao et al., 2014). Following the same reasoning, the morpheme-based fluency task may be even more demanding compared with the standard fluency tasks, as it requires more selective inhibition of inappropriate responses, e.g., semantic associations or inflections of previously provided words. The current data suggest that the morpheme-based tasks (both root-based and pattern-based) were more difficult than the standard fluency tasks, although a direct comparison is unwarranted due to the different administering procedures (see Methods). The extent to which morpheme-based fluency relies on executive functions was beyond the scope of the current investigation and remains to be directly tested in future research.

In the current study, neurocognitive correlations were found between properties of white matter tracts and root-based fluency. In contrast, a correlation with pattern-based fluency could only be found when we included FR in our analyses, and its location was inconsistent across the two analysis procedures. This discrepancy between the two morphemes is in accordance with behavioral studies in Hebrew, which report that pattern-based priming effects are weaker and less consistent than effects obtained using the root (Deutsch et al., 2005; Frost et al., 2000, 1997). Interestingly, pattern effects could be restored using behavioral paradigms that intentionally abolish the more dominant effect of the root (Deutsch et al., 2018). This attests to the higher salience of the root compared with the pattern, which results from the different roles these morphemes play in Hebrew word structure. The consonantal root carries the core meaning of the word, while the pattern, which consists mostly of vowels, provides its phonological template. Indeed, we found that pattern-based fluency was strongly correlated with letter-based fluency, suggesting that these tasks rely on similar phonological processes. Taken together, the body of research in Hebrew suggests that while both root and pattern play a part in lexical access (see, for example, Deutsch and Kuperman, 2019), the pattern morpheme conveys less consistent mapping to meaning, and is more sensitive to experimental design. Whether more sensitive behavioral paradigms would reveal associations between pattern-related effects and white matter tracts remains a question for future research.

### 4.3. Limitations

The correlational nature of the study does not allow us to provide a mechanistic explanation for the involvement of white matter tracts in morpheme-based fluency or to determine whether different tracts support different aspects of performance in this task. Studies involving direct training of specific morphological abilities may be required to draw conclusions about the causal role of these tracts in cognition. While we interpret the findings in the ventral tracts as confirming their contribution to amodal morpheme-based lexical access, we cannot rule out the possibility that these correlations arise as a result of other cognitive processes involved in this task. For example, participants could use orthographic strategies and visual imagery while searching for morphologically-related words. In this case, the effects observed in the ventral tracts may reflect their well-known involvement in orthographic processing (Banfi et al., 2019; Horowitz-Kraus et al., 2014; Huber et al., 2018; Ozernov-Palchik et al., 2019; Vandermosten et al., 2012; Yeatman et al., 2012a). Although the current data do no not allow to preclude this alternative explanation of the results, it seems less plausible given that morpheme-based fluency was not correlated with reading ability. In addition, the morphological effects in the ventral pathways remained significant when controlling for letter-based fluency, which may also involve orthographic strategies (Friedman et al., 1998). The broader question of whether morpheme-based fluency is related to reading ability deserves additional research, given prior indications that dyslexics generally perform poorly in this task (Ben-Dror et al., 1995; Leikin and Even Zur, 2006; Su et al., 2018). Critically, the measure used here to evaluate reading was an oral single word reading test, which may not capture the most relevant variance to address this question. Future studies may incorporate more carefully designed reading evaluation tasks (for example, using silent reading or text comprehension), and include participants with a larger range of reading abilities in the sample. Lastly, the morpheme-based fluency tasks were not directly comparable to the standard category- and letter-based fluency tasks, because more items were administered in the morpheme-based tasks, and participants were given less time to respond to each of these items (see Methods). This difference may not be critical, because there are, by definition, far fewer words that share a specific morphemes than words that begin with the same letter or belong to the semantic categories used here. Indeed, participants reached their maximum capacity to come up with new morphologically-related words before approaching the time limit. While the comparison between the tasks was not the focus of the study, the different procedures make it difficult to interpret the current results in the context of previous literature on verbal fluency.

### 4.4. Summary and conclusions

The results of the current study show that morpheme-based fluency is associated with both ventral and dorsal white matter tracts, bilaterally. The same ventral tracts were previously shown to be involved in morphological processing in visual word recognition across both English and Hebrew, languages that markedly differ in morphological structure and writing system (Yablonski et al., 2019; Yablonski and Ben-Shachar, 2020). The current results further expand these findings to the production domain, suggesting that the ventral tracts are involved in morphological processing beyond reading, thereby generalizing previous findings across task, input modality and response modality. These findings support a view of morphemes as amodal entities which govern lexical access, by mapping either written or spoken word forms to the semantic system. Moreover, the involvement of the bilateral FAT in morpheme-based fluency provides new evidence supporting the importance of this tract for speech production in typical adults, extending the clinically-focused verbal fluency literature. Together, the results of the current study emphasize that neurocognitive correlations reflect the linguistic construct under investigation but are also largely modulated by task selection. Lastly, this study makes an important contribution to the literature by incorporating a novel measurement of fiber density, FR. Using FR we were able to uncover additional correlations with morpheme-based fluency that could not be detected using standard tensor-derived metrics. Although more research will be required to elucidate the biological underpinnings of FR, our findings clearly demonstrate the added value of including novel structural measurements when investigating neurocognitive associations within white matter pathways.

## Acknowledgements

This study was conducted as part of Maya Yablonski’s doctoral dissertation, carried under the supervision of Prof. Michal Ben-Shachar at the Gonda Multidisciplinary Brain Research Center, Bar-Ilan University. This study was supported by the Israel Science Foundation (ISF grant #1083/17), the US-Israel Binational Science Foundation (BSF grant #2011314) and the center for research excellence in cognitive science (I-CORE Program 51/11). Maya Yablonski was partly supported by the Wolf Foundation. We are grateful to Yaniv Assaf for guidance in implementing the CHARMED protocol and analysis pipeline. We thank Vered Kronfeld-Duenias for assistance in analysis of CHARMED data. We thank Daniel Barazany and the team at the Strauss Center for Computational Neuroimaging for their assistance in protocol setup and MRI scanning. We thank Kathleen Rastle and JSH Taylor for helpful discussions.

