## SupplementaryMaterial for "A general role for ventral white matter pathways in morphological processing: going beyond reading"

### Supplemental text

#### Methods

#### Deterministic tractography

Data preprocessing, including tensor fitting, was conducted using the same steps as in the main analysis (See *Methods, 2.3.2.1-2*). Then, deterministic whole-brain tractography was carried out using AFQ software, using a Streamlines Tractography (STT) algorithm, with a 4th-order Runge–Kutta path integration method and 1 mm step size (Mori et al., 1999; Pierpaoli et al., 2001). Tracking was initiated within a white matter mask containing all brain voxels with FA > 0.2. The tracking algorithm was halted when it reached voxels with FA < 0.2 or an angle greater than 30° between the last and the next step direction. Minimum and maximum streamline length were set to 20mm and 250mm, respectively. The resulting tracts were cleaned automatically, using a statistical outlier rejection algorithm that removed streamlines that were more than 4 standard deviations longer than the mean tract length, or that deviate by more than 5 standard deviations in distance from the tract core. This process was iterated 5 times for each tract (see (Yeatman et al., 2012) for details regarding the automatic segmentation method).

#### Results

The twelve tracts of interest were successfully detected in the majority of participants, with the following exceptions: the right AF-ft could not be segmented in 5/45 participants, the left FAT in 3/45 and right FAT in 4/45. In one participant the segmented FAT consistent of spurious streamlines that did not fit the tract anatomical trajectory, and was excluded from analyses bilaterally. Shapiro-Wilk test confirmed that the mean tract FA and MD in each of the tracts were normally distributed following outlier exclusion (see *Methods*). In total, 0-2 subjects were excluded from subsequent analyses of each tract (actual sample sizes included in correlation analyses for each tract are reported in Tables S4, S5 and S8 for FA, MD and FR, respectively).

Supplemental Tables S6, S7, S9 and Figures S4-S5, S8-S10 report the results of the tensor-deterministic analysis in parallel to the results of the probabilistic pipeline, reported in the main text. The other Figures provide further analyses of the diffusion metrics in the current dataset: Figures S1-S3 delineate the comparison between diffusion metrics extracted from tracts reconstructed using the probabilistic pipeline with those reconstructed using the tensor-deterministic approach. Figures S11-S13 demonstrate how FR covaries with the traditional tensor-derived metrics. Each Supplemental Table and Figure is referred to from the main text.

**Supplemental Figures**

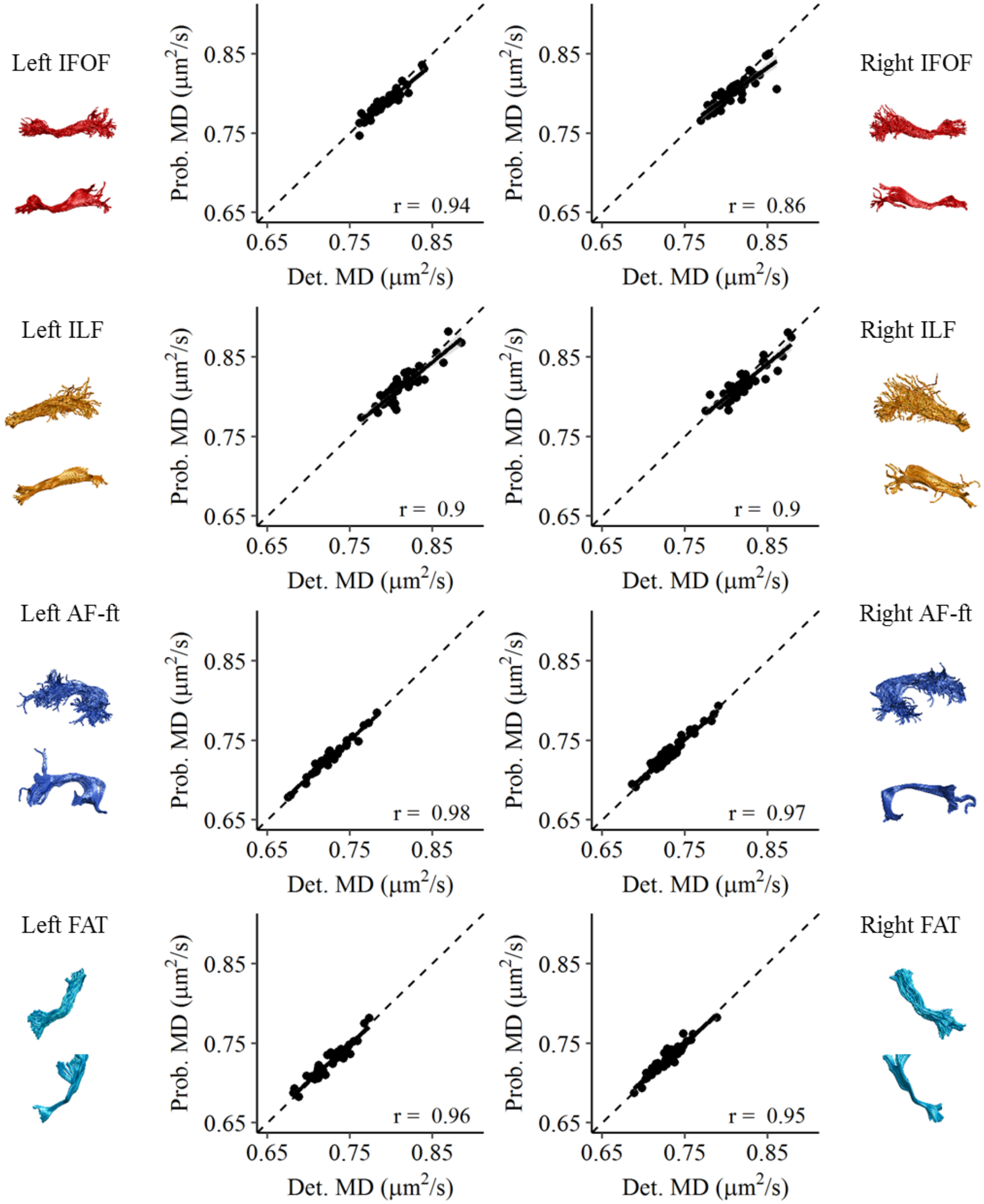


**Figure S1.** Consistency of mean MD extracted from tracts using the two tractography pipelined: Det. - deterministic tractography with tensor modeling and Prob. - probabilistic tractography with CSD modeling. MD values were extracted from tracts reconstructed using the two pipelines and averaged across the 100 nodes. R values denote Pearson's correlation coefficient. Black lines represent the best linear fit, surrounded by the 95% confidence interval (shaded area). Dashed lines represent y=x.

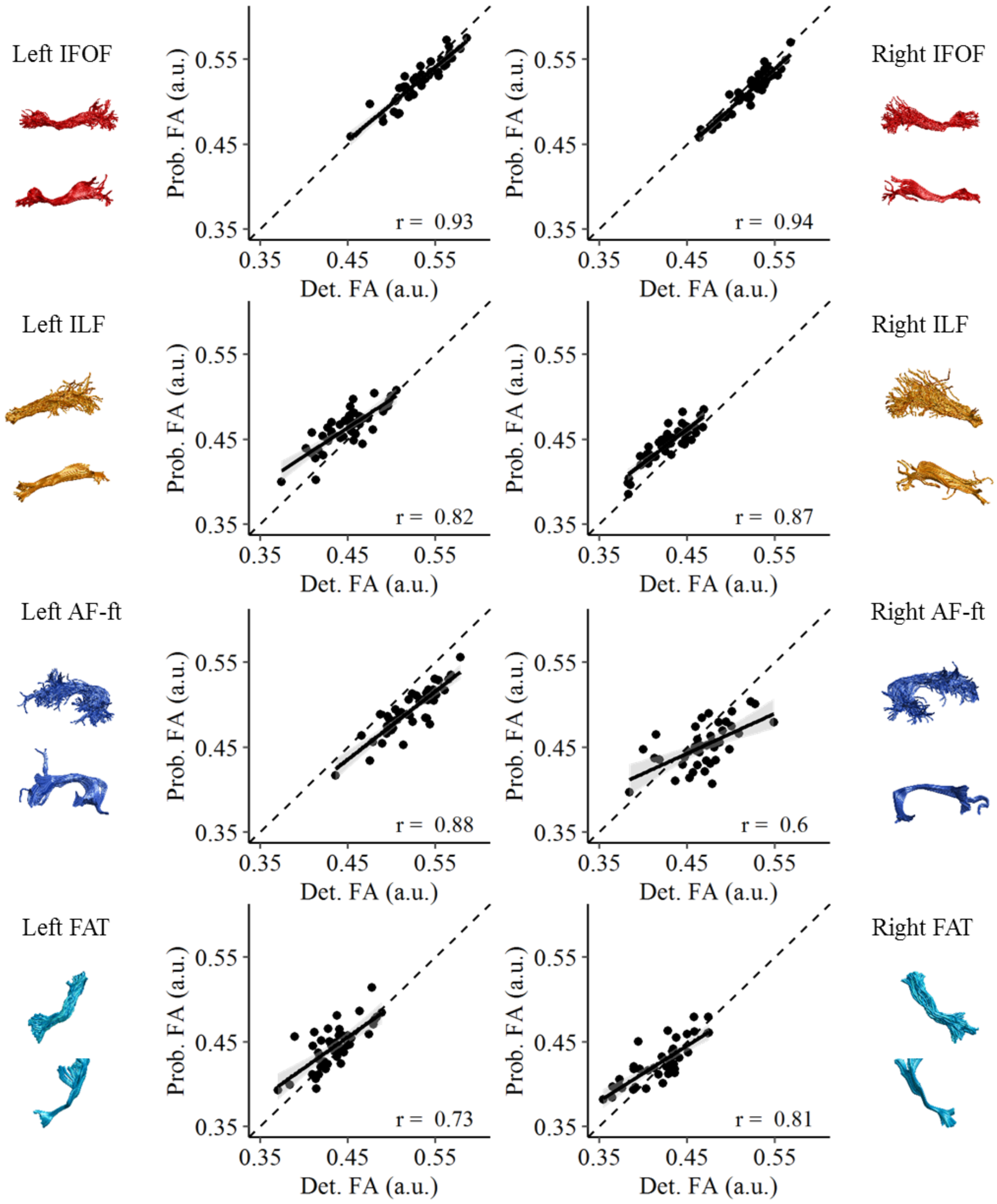


**Figure S2.** Consistency of mean FA extracted from tracts using the two tractography pipelined: Det. - deterministic tractography with tensor modeling and Prob. - probabilistic tractography with CSD modeling. FA values were extracted from tracts reconstructed using the two pipelines and averaged across the 100 nodes. R values denote Pearson's correlation coefficient. Black lines represent the best linear fit, surrounded by the 95% confidence interval (shaded area). Dashed lines represent y=x.

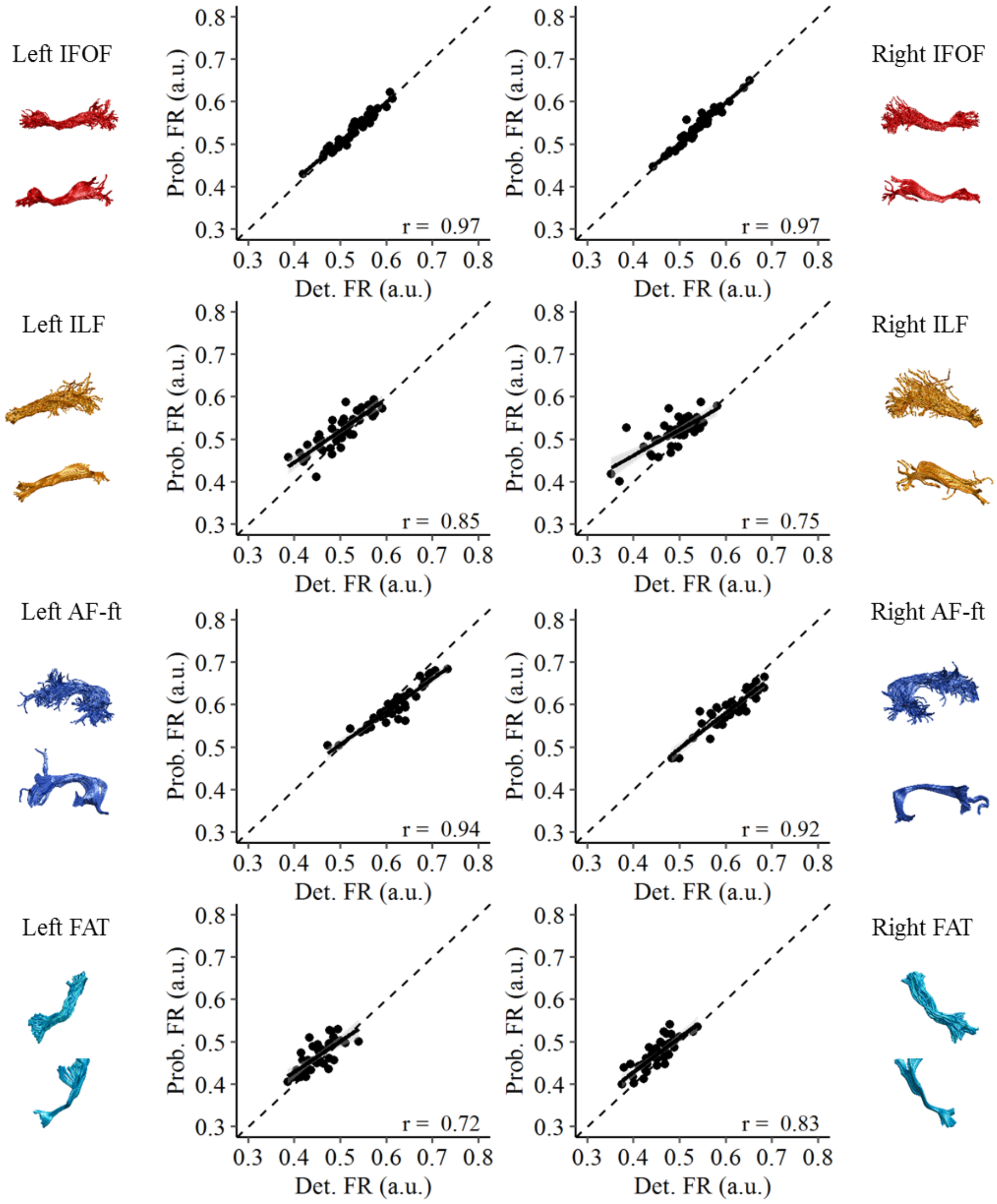


**Figure S3.** Consistency of mean FR extracted from tracts using the two tractography pipelined: Det. - deterministic tractography with tensor modeling and Prob. - probabilistic tractography with CSD modeling. FR values were extracted from tracts reconstructed using the two pipelines and averaged across the 100 nodes. R values denote Pearson's correlation coefficient. Black lines represent the best linear fit, surrounded by the 95% confidence interval (shaded area). Dashed lines represent y=x.

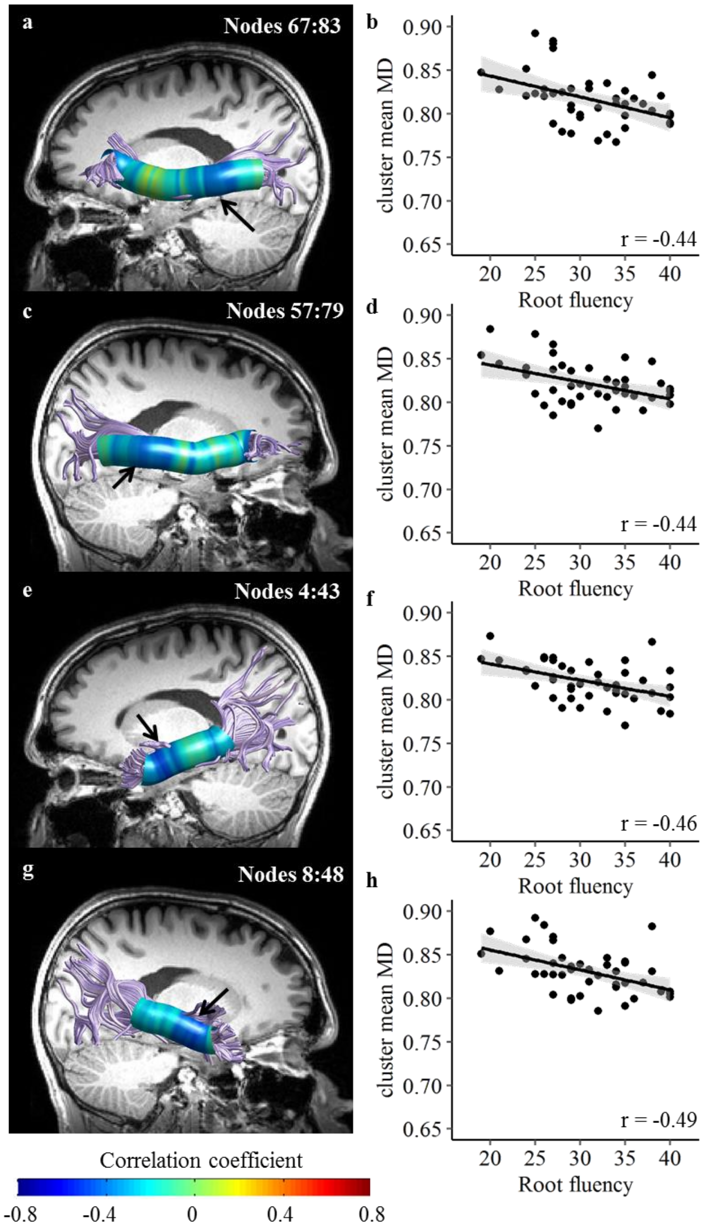


**Figure S4. Root-based fluency is negatively correlated with MD in bilateral ventral tracts.** This figure parallels Figure 3 in the main manuscript, with tracts reconstructed using deterministic tractography algorithm and tensor-based modeling. (a, c, e, g) Pearson's correlation coefficients are visualized in 100 nodes along the left IFOF (a), right IFOF (c), left ILF (e) and right ILF (g). Black arrows denote the location of significant clusters after family-wise error correction across the 100 nodes. (b, d, f, h) Scatter plots showing the association between root-based fluency (number of words) and the mean MD in the significant cluster of nodes, in left IFOF (b), right IFOF (d), left ILF (f) and right ILF (h). Black lines represent the best linear fit, surrounded by the 95% confidence interval (shaded area). These scatter plots are shown for visualization purposes, significance is calculated along the trajectory of the tracts. IFOF- inferior fronto-occipital fasciculus. ILF- inferior longitudinal fasciculus.

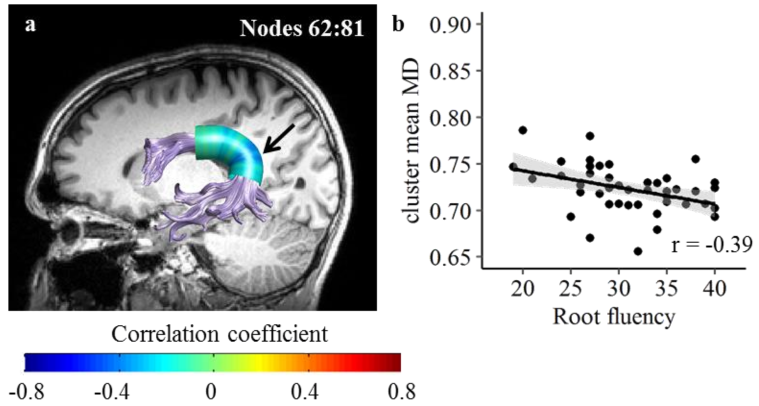


**Figure S5. Root based fluency is negatively correlated with MD in the left AF-ft** (a) Pearson's correlation coefficients are visualized in 100 nodes along the left AF-ft. Black arrows denote the location of significant clusters after family-wise error correction across the 100 nodes. (b) Scatter plots showing the association between root-based fluency (number of words) and the mean MD in the significant cluster of nodes. Black lines represent the best linear fit, surrounded by the 95% confidence interval (shaded area). The scatter plot is shown for visualization purposes, significance is calculated along the trajectory of the tracts. AF-ft – Arcuate Fasciculus, fronto-temporal segment

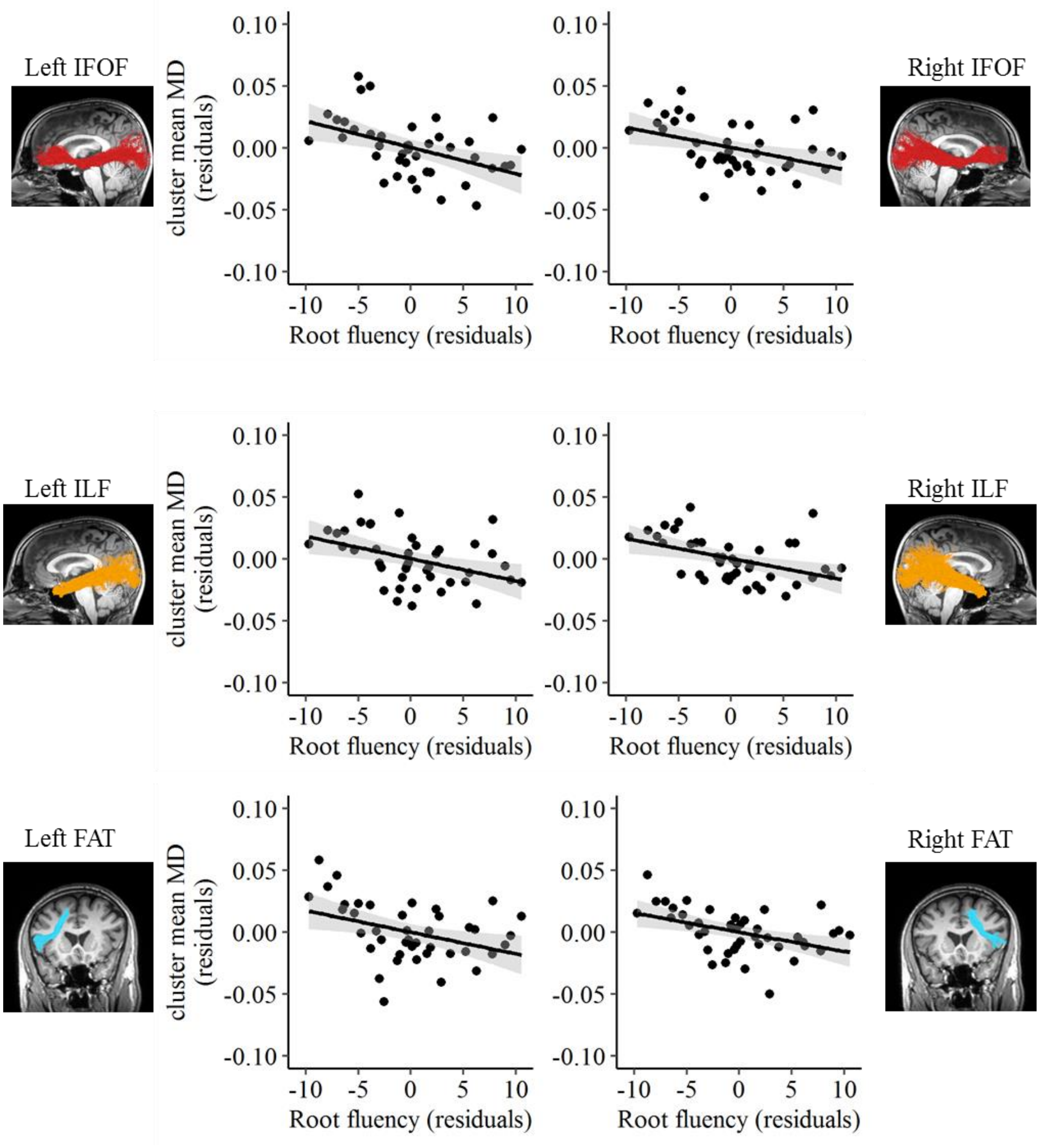


**Figure S6. Correlations between MD and root fluency remain significant after controlling for category-based fluency, letter- based fluency, and age.** Partial correlations are visualized as scatter plots between root fluency residuals and MD residuals, regressed on category-based fluency, letter-based fluency and age. These scatterplots parallel those presented in Figure 3 and Figure 5 in the main text. Root-based fluency was a significant predictor of MD in all clusters, while the contribution of the other predictors was non-significant ($p > 0.1$; except for the right ILF where the contribution of letter-based fluency was near significance, $p = 0.0504$).

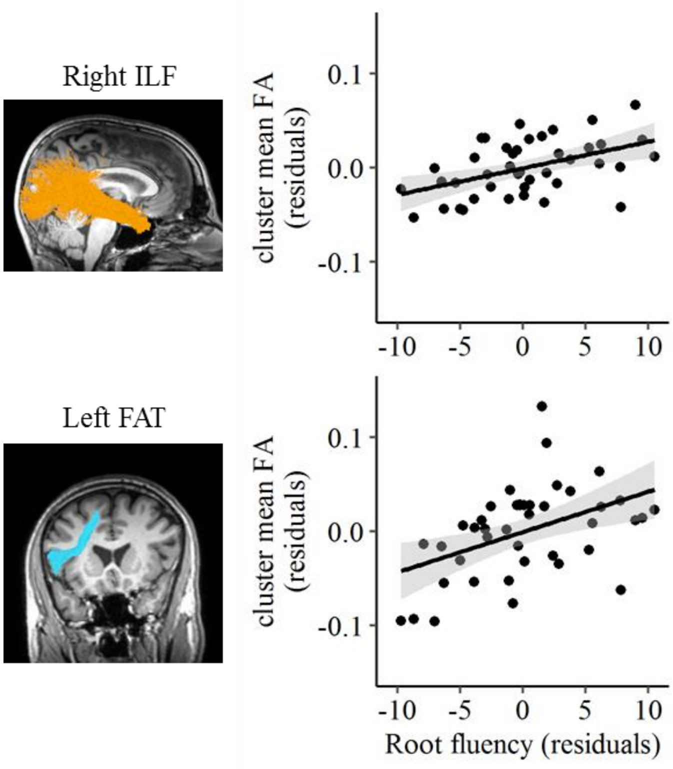


**Figure S7. Correlations between FA and root fluency remain significant after controlling for category-based fluency, letter- based fluency, and age.** Partial correlations are visualized as scatter plots between root fluency residuals and FA residuals, regressed on category-based fluency, letter-based fluency and age. These scatterplots parallel those presented in Figure 4 in the main text. Root-based fluency was a significant predictor of FA in all clusters, while the contribution of the other predictors was non-significant ($p > 0.1$).

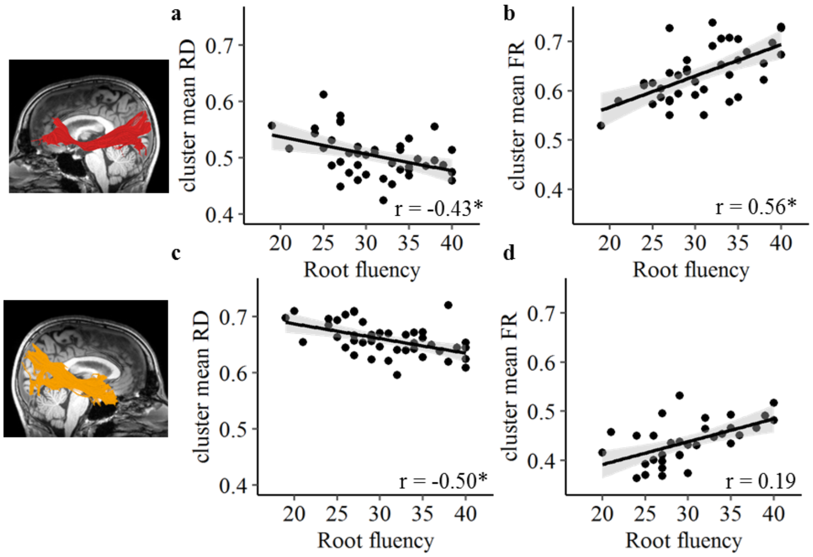


**Figure S8. Root-based fluency is correlated with RD in several tracts, yet correlations with FR are found only in the left IFOF.** Scatter plots show the association between root-based fluency and mean radial diffusivity (RD; panels a, c) or mean diffusivity in the restricted compartment (FR; panels b, d) within clusters showing a significant correlation between RD and root-based fluency (see Table S8 for a full report of the correlations with RD and FR in all the significant clusters). This Figure parallels Figure 6 in the main manuscript, for deterministic tracts. Tractograms on the left demonstrate the relevant pathways in a single participant. $*p < 0.05$, FDR corrected for 15 multiple comparisons (3 diffusion measurements x 5 clusters that showed significant correlations with root-based fluency).

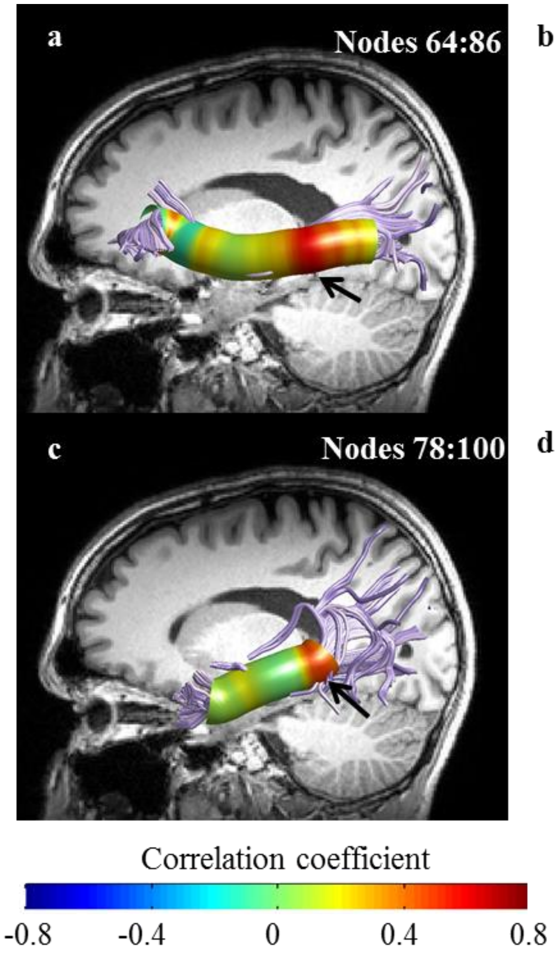


**Figure S9. Analysis of FR along the tracts reveals selective correlations with ventral tracts in the left hemisphere.** This figure parallels Figure 7 in the main manuscript, with tracts reconstructed using deterministic tractography algorithm and tensor-based modeling. Root based fluency is positively correlated with FR in significant clusters in the left IFOF (a-b) and left ILF (c-d). No significant clusters were found in the right hemisphere tracts that showed significant FA or MD effects. Black lines represent the best linear fit, surrounded by the 95% confidence interval (shaded area). These scatter plots are shown for visualization purposes, significance is calculated along the trajectory of the tracts. IFOF- inferior fronto-occipital fasciculus. ILF- inferior longitudinal fasciculus.

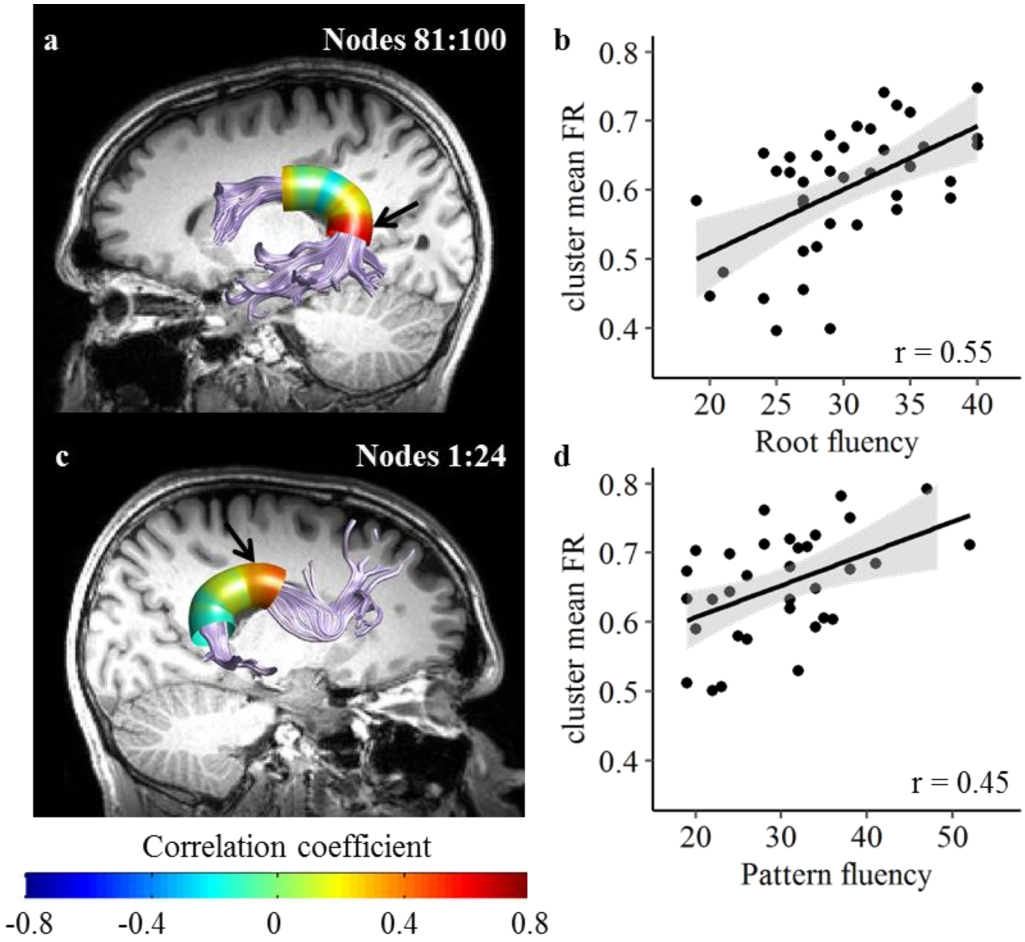


**Figure S10. Analysis of FR along the tracts uncovers correlations that were not detected using traditional diffusion metrics.** This figure parallels Figure 8 in the main manuscript, with tracts reconstructed using deterministic tractography algorithm and tensor-based modeling. Root based fluency is correlated with FR in a significant cluster in the left AF-ft (a-b). Pattern based fluency is correlated with FR in a significant cluster in the right AF-ft (c-d). Black lines represent the best linear fit, surrounded by the 95% confidence interval (shaded area). These scatter plots are shown for visualization purposes, significance is calculated along the trajectory of the tracts. AF-ft- fronto-temporal segment of the arcuate fasciculus.

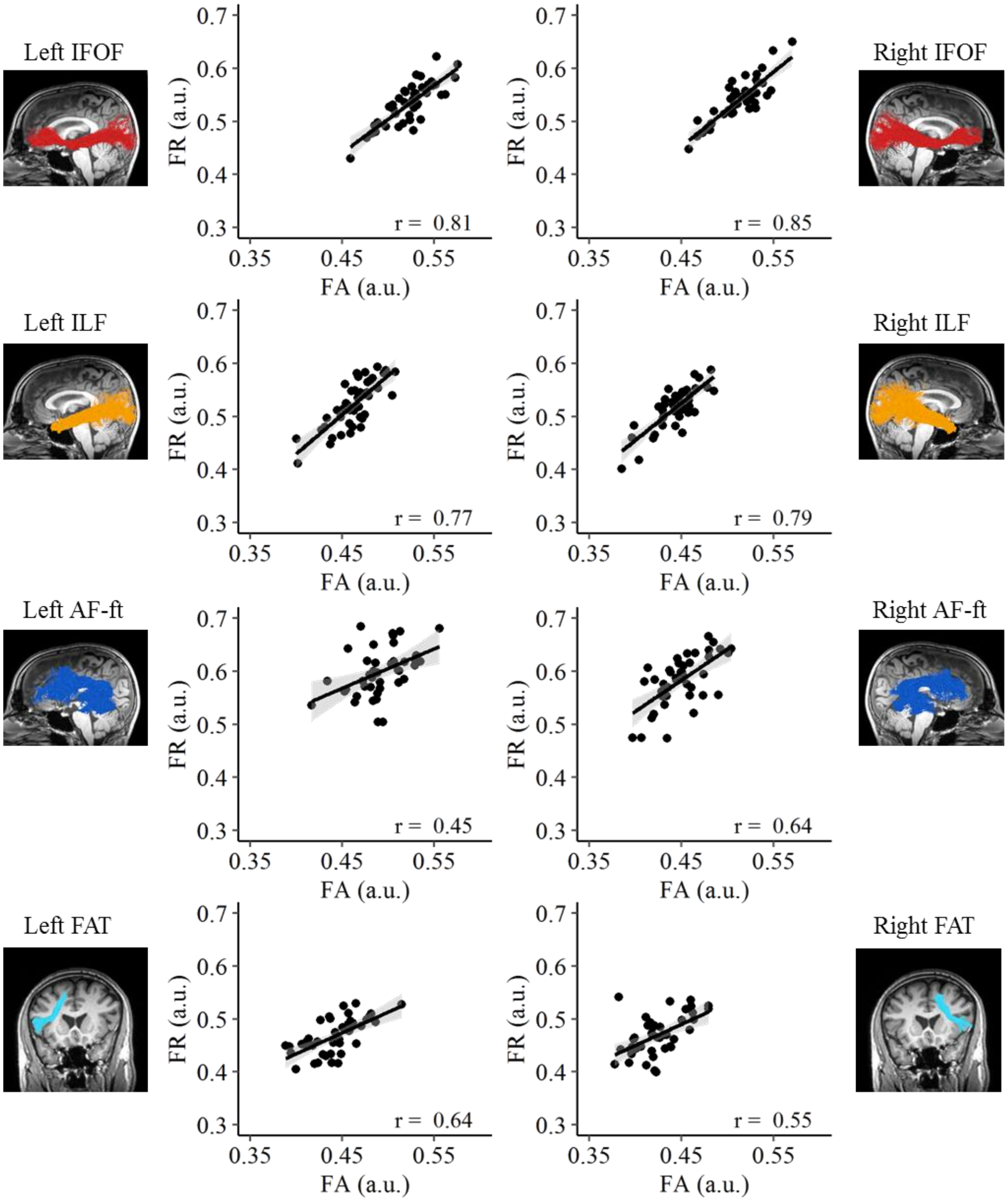


**Figure S11. Correlations between mean FR and mean FA vary across different tracts.** FR and FA were extracted from tracts reconstructed using probabilistic tractography and averaged across the 100 nodes. R values denote Pearson's correlation coefficient. Black lines represent the best linear fit, surrounded by the 95% confidence interval (shaded area).

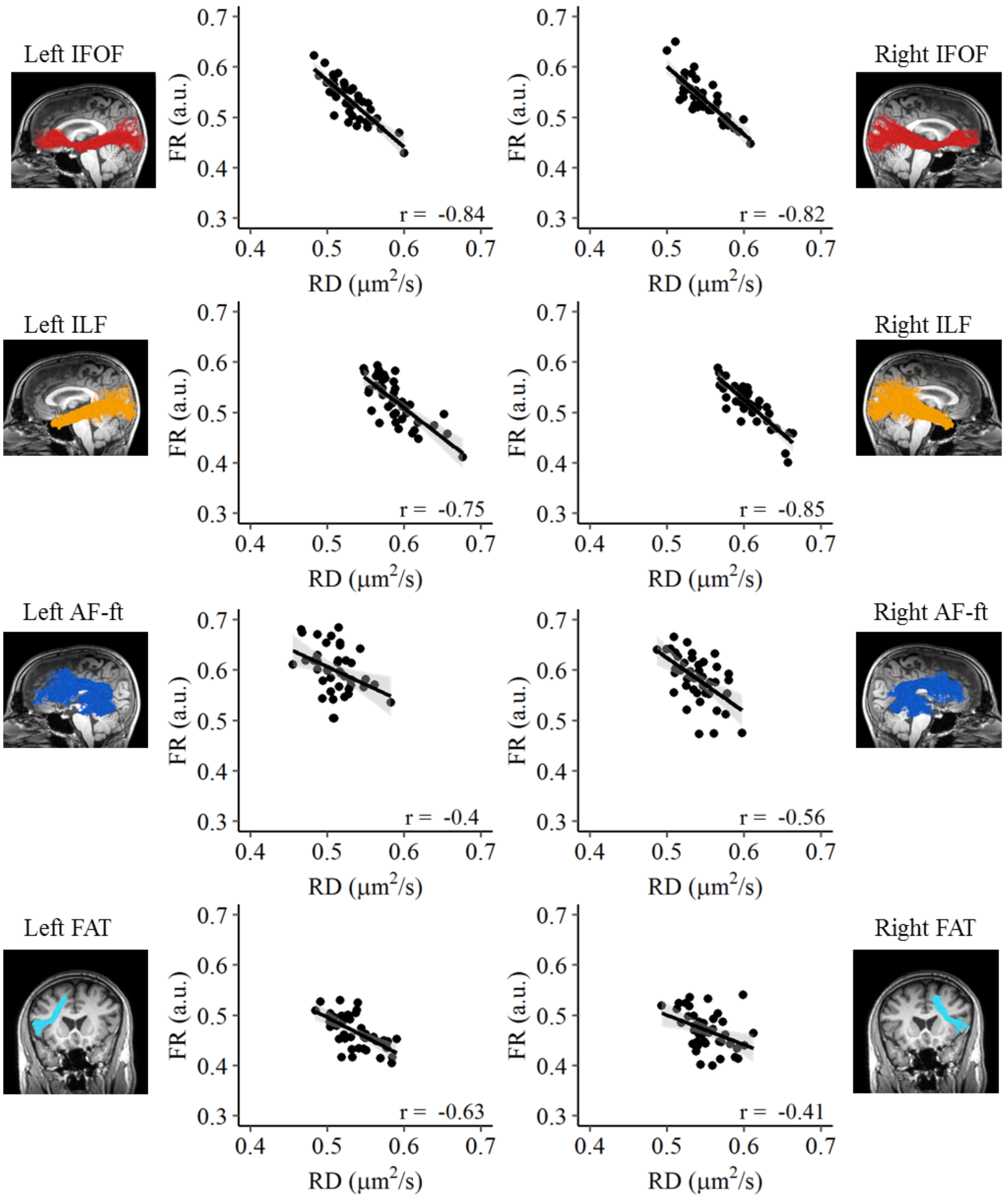


**Figure S12.** Correlations between mean FR and mean RD vary across different tracts. FR and RD were extracted from tracts reconstructed using probabilistic tractography and averaged across the 100 nodes. R values denote Pearson's correlation coefficient. Black lines represent the best linear fit, surrounded by the 95% confidence interval (shaded area).

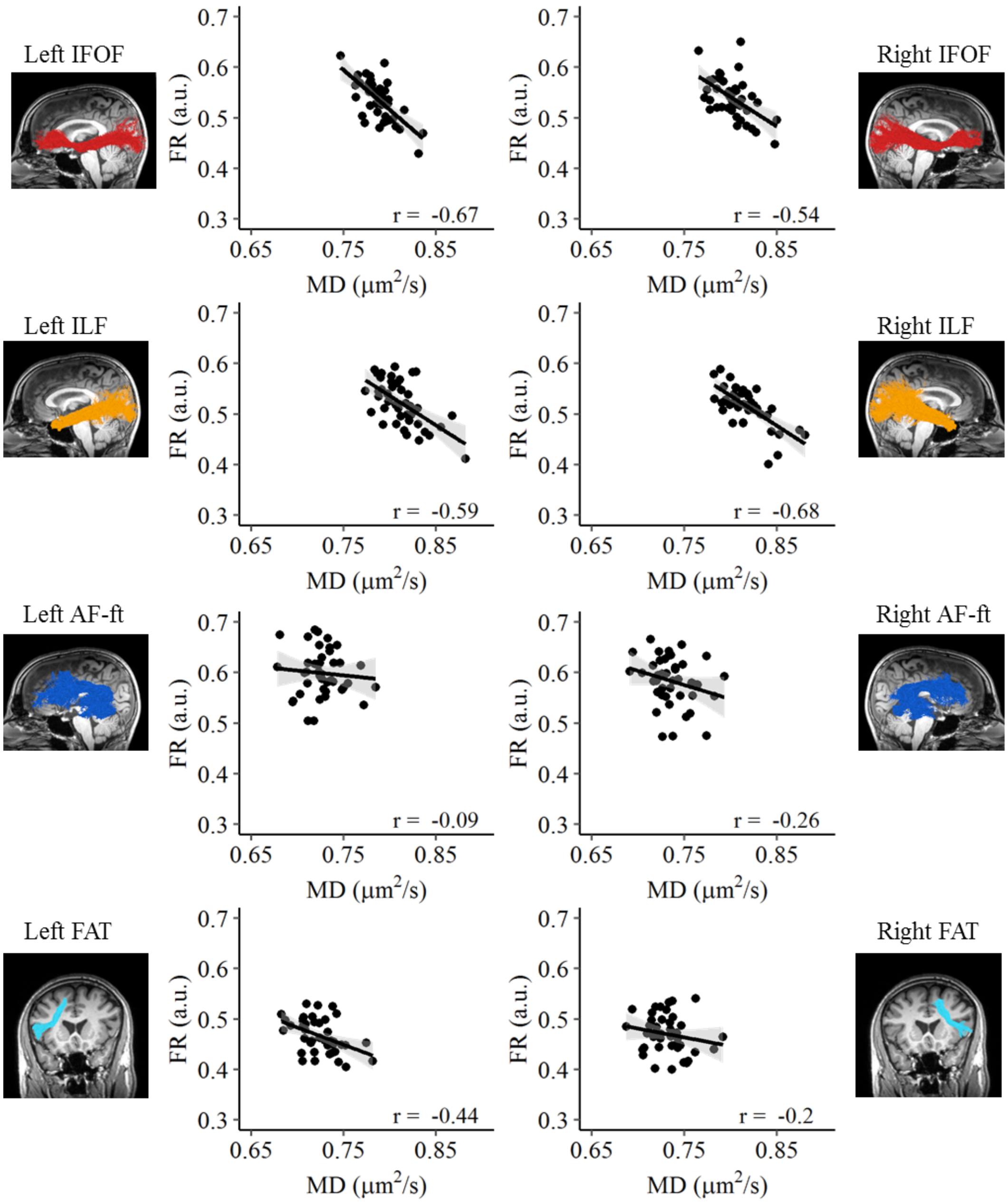


**Figure S13.** Correlations between mean FR and mean MD vary across different tracts. FR and MD were extracted from tracts reconstructed using probabilistic tractography and averaged across the 100 nodes. R values denote Pearson's correlation coefficient. Black lines represent the best linear fit, surrounded by the 95% confidence interval (shaded area).

### Supplemental Tables

| N=45 | Mean | SD | Range |
|---|---|---|---|
| **Fluency tasks** | | | |
| Root | 30.89 | 5.54 | [19, 40] |
| Pattern | 29.70 | 9.44 | [15, 54] |
| Letter (Z-score) | 0.16 | 1.03 | [-2.41, 2.37] |
| Category (Z-score) | 0.65 | 1.17 | [-1.40, 3.52] |
| **Background measures** | | | |
| Gender | 16M/29F | | |
| Age (years) | 26.45 | 3.72 | [20.23, 34.87] |
| Education (years) | 14.56 | 2.13 | [12, 20] |
| Handedness (laterality quotient) | 96.89 | 5.90 | [80,100] |
| Single word reading (#words) | 99.07 | 20.54 | [62,156] |

**Table S1. Performance on fluency tasks and background measures.**

Handedness scores are based on the Edinburgh handedness inventory (Oldfield, 1971), 100 indicates full right handedness, -100 indicates full left handedness. Single word reading- number of words correctly read aloud in 1 minute (Shatil, 1997).

| Root-based fluency | | | Pattern-based fluency | | |
|---|---|---|---|---|---|
| miSGeRet | a frame | מסגרת | MeLeX | a king | מלך |
| KaTaV | wrote | כתב | ŠMiRa | guarding | שמירה |
| SaDRan | an usher | סדרן | XaRuC | hard-working | חרוץ |
| maFTeaX | a key | מפתח | maCLeMa | a camera | מצלמה |
| maXŠaVa | a thought | מחשבה | haFTaʔa | a surprise | הפתעה |

**Table S2. Stimuli used as targets for the morpheme-based fluency task.** These stimuli are a subset of the stimuli used in (Leikin and Even Zur, 2006). Capital letters denote root consonants. All targets were read aloud by the experimenter and no part of the task involved reading.

| | Root | Pattern | Letter | Category | Age | Education |
|---|---|---|---|---|---|---|
| Root | | | | | | |
| Pattern | 0.38 | | | | | |
| Letter | 0.33 | **0.60*** | | | | |
| Category | 0.38 | 0.36 | **0.53*** | | | |
| Age | 0.06 | 0.24 | 0.36 | -0.09 | | |
| Education | 0.27 | 0.17 | 0.39 | -0.25 | **0.67*** | |
| Word Reading | 0.22 | **0.45*** | **0.46*** | **0.42*** | 0.10 | 0.20 |

**Table S3. Correlations between the fluency tasks and background measures.** Shown are two-tailed Pearson's correlation coefficients between the behavioral measures collected in the sample (N=44). Values in bold with an asterisk are significant by $p < 0.05$, FDR corrected for 21 correlations.

| | **Probabilistic** | | | **Deterministic** | | |
|---|---|---|---|---|---|---|
| | **r** | **p** | **N** | **r** | **p** | **N** |
| Left IFOF | 0.39 | 0.010 | 43 | 0.38 | 0.013 | 43 |
| Right IFOF | 0.40 | 0.008 | 44 | 0.44 | **0.003*** | 44 |
| Left ILF | 0.32 | 0.036 | 42 | 0.37 | 0.013 | 43 |
| Right ILF | 0.22 | 0.163 | 43 | 0.27 | 0.082 | 44 |
| Left AF-ft | 0.25 | 0.108 | 43 | 0.21 | 0.169 | 43 |
| Right AF-ft | 0.21 | 0.162 | 44 | 0.17 | 0.307 | 38 |
| Left UF | 0.31 | 0.041 | 43 | 0.13 | 0.388 | 44 |
| Right UF | 0.19 | 0.219 | 43 | 0.10 | 0.531 | 43 |
| Left FAT | 0.34 | 0.027 | 43 | 0.24 | 0.143 | 39 |
| Right FAT | 0.23 | 0.132 | 44 | 0.05 | 0.740 | 39 |
| Left AF-fp | 0.11 | 0.487 | 44 | 0.08 | 0.610 | 44 |
| Right AF-fp | 0.15 | 0.346 | 43 | -0.09 | 0.551 | 43 |

**Table S4. Correlations between root-based fluency and mean tract FA, in tracts reconstructed using either a probabilistic or deterministic tracking algorithm.** N represents the actual sample size included in the analyses of each tract after outlier exclusion. *$p < 0.05$, FDR corrected for the 12 tracts of interest.

| | Probabilistic | | | Deterministic | | |
|---|---|---|---|---|---|---|
| | **r** | **p** | **N** | **r** | **p** | **N** |
| Left IFOF | -0.32 | 0.041 | 42 | -0.40 | **0.008*** | 43 |
| Right IFOF | -0.25 | 0.104 | 42 | -0.42 | **0.006*** | 43 |
| Left ILF | -0.38 | 0.012 | 43 | -0.38 | **0.012*** | 43 |
| Right ILF | -0.33 | 0.035 | 42 | -0.41 | **0.006*** | 44 |
| Left AF-ft | -0.26 | 0.095 | 43 | -0.29 | 0.061 | 43 |
| Right AF-ft | -0.16 | 0.306 | 43 | -0.25 | 0.118 | 39 |
| Left UF | -0.30 | 0.053 | 42 | -0.29 | 0.062 | 42 |
| Right UF | -0.07 | 0.675 | 42 | -0.04 | 0.808 | 42 |
| Left FAT | -0.33 | 0.032 | 43 | -0.23 | 0.158 | 39 |
| Right FAT | -0.41 | 0.007 | 43 | -0.24 | 0.144 | 38 |
| Left AF-fp | -0.18 | 0.254 | 43 | -0.26 | 0.096 | 43 |
| Right AF-fp | -0.15 | 0.337 | 43 | -0.14 | 0.379 | 44 |

**Table S5. Correlations between root-based fluency and mean tract MD, in tracts reconstructed using either a probabilistic or deterministic tracking algorithm.** N represents the actual sample size included in the analyses of each tract after outlier exclusion. *$p < 0.05$, FDR corrected for the 12 tracts of interest.

| | cluster location (Nodes) | r | p | CI (95%) | Regression Full model $R^2$(%) | Regression Root fluency p |
|---|---|---|---|---|---|---|
| Left IFOF | 67 – 83 | -0.44 | 0.003* | [-0.58,-0.21] | 21% | 0.007§ |
| Right IFOF | 57 – 79 | -0.44 | 0.003* | [-0.65, -0.16] | 20% | 0.010§ |
| Left ILF | 4 – 43 | -0.46 | 0.002* | [-0.68, -0.10] | 32% | 0.002§ |
| Right ILF | 8 – 48 | -0.49 | 0.001* | [-0.66, -0.13] | 24% | 0.002§ |
| Left AF-ft | 62 – 81 | -0.39 | 0.010* | [-0.62, -0.10] | 23% | 0.003§ |

**Table S6. Root-based fluency is correlated with MD in ventral and dorsal tracts reconstructed using deterministic tractography.** This table parallels Table 1 from the main text, with values extracted from deterministic tracts. Reported are clusters of nodes showing significant Pearson's correlations with Root-based fluency, family-wise error corrected for 100 nodes. Significant clusters were followed up with multiple regression models predicting cluster mean MD from root-based fluency, category-based fluency, letter-based fluency and age. We report the R squared of each regression model and the significance level of the root fluency predictor. The contribution of the additional predictor variables was non-significant in all the models tested ($p > 0.05$), except for the left ILF, where letter-based fluency made a significant contribution ($p = 0.019$). *$p < 0.05$, FDR corrected for 5 clusters. § $p < 0.05$ for the root-based fluency predictor within each regression model

| | cluster location (Nodes) | AD | p | RD | p | FR | p |
|---|---|---|---|---|---|---|---|
| Left IFOF | 67 – 83 | -0.13 | 0.421 | -0.43 | **0.004*** | 0.56 | **$2x10^{-4}$*** |
| Right IFOF | 57 – 79 | -0.33 | 0.033 | -0.28 | 0.067 | 0.25 | 0.125 |
| Left ILF | 4 – 43 | -0.19 | 0.225 | -0.37 | 0.014 | 0.11 | 0.505 |
| Right ILF | 8 – 48 | -0.21 | 0.162 | -0.50 | **0.001*** | 0.19 | 0.222 |
| Left AF-ft | 62 – 81 | -0.20 | 0.197 | -0.18 | 0.245 | 0.11 | 0.480 |

**Table S7. Root-based fluency is correlated with FR in the left IFOF.** This table parallels Table 2 from the main text, with values extracted from deterministic tracts. For each significant cluster we extracted the mean value of axial diffusivity (AD), radial diffusivity (RD) and the restricted diffusion fraction from the CHARMED model (FR), an index of fiber density. Although clusters in several tracts show a significant correlation with RD, correlations with FR are found in the left IFOF alone. Reported are Pearson's correlation coefficients between each diffusion component and root fluency. *$p < 0.05$, FDR corrected for 15 comparisons (5 clusters * 3 measurements)

| | Probabilistic | | | Deterministic | | |
|---|---|---|---|---|---|---|
| | **r** | **p** | **N** | **r** | **p** | **N** |
| Left IFOF | 0.42 | 0.007 | 40 | 0.41 | 0.009 | 40 |
| Right IFOF | 0.32 | 0.045 | 40 | 0.36 | 0.024 | 40 |
| Left ILF | 0.31 | 0.047 | 41 | 0.22 | 0.163 | 41 |
| Right ILF | 0.38 | 0.017 | 39 | 0.28 | 0.082 | 40 |
| Left AF-ft | 0.17 | 0.296 | 41 | 0.28 | 0.086 | 40 |
| Right AF-ft | 0.13 | 0.401 | 41 | 0.17 | 0.327 | 36 |
| Left UF | 0.13 | 0.424 | 41 | -0.01 | 0.963 | 41 |
| Right UF | 0.18 | 0.273 | 40 | 0.08 | 0.613 | 41 |
| Left FAT | 0.26 | 0.094 | 41 | 0.16 | 0.352 | 35 |
| Right FAT | -0.01 | 0.932 | 41 | -0.33 | 0.043 | 37 |
| Left AF-fp | -0.06 | 0.689 | 41 | 0.04 | 0.800 | 41 |
| Right AF-fp | 0.18 | 0.267 | 41 | 0.10 | 0.548 | 41 |

**Table S8. Correlations between root-based fluency and mean tract FR, in tracts reconstructed using either a probabilistic or deterministic tracking algorithm.** None of the correlations was significant at a significance level of 0.05 corrected for the 12 tracts of interest.

| | Cluster location (Nodes) | r | p | CI (95%) | Regression Full model $R^2$(%) | Regression Morpheme-based fluency p |
|---|---|---|---|---|---|---|
| Root-based fluency | | | | | | |
| Left IFOF | 64 – 86 | 0.54 | $3x10^{-4}$* | [0.29, 0.73] | 36 | $2x10^{-4§}$ |
| Left ILF | 78 – 100 | 0.46 | 0.002* | [0.21, 0.66] | 26 | $0.002^{§}$ |
| Left AF-ft | 81 – 100 | 0.55 | $2x10^{-4}$* | [0.35, 0.71] | 39 | $0.001^{§}$ |
| Pattern-based fluency | | | | | | |
| Right AF-ft | 1 – 24 | 0.45 | 0.006* | [0.18,0.65] | 33 | $0.042^{§}$ |

**Table S9. Diffusivity in the restricted compartment (FR) is correlated with root-based fluency and pattern-based fluency.** This table parallels Table 3 in the main text, with values extracted from deterministic tracts. Reported are clusters of nodes showing significant Pearson's correlations between FR and Root-based fluency or Pattern-based fluency, family-wise error corrected for 100 nodes. Significant clusters were followed up with multiple regression models predicting cluster mean FR from root-based or pattern-based fluency, as well as category-based fluency, letter-based fluency and age. We report the R squared of each regression model and the significance level of the predictor of interest, root or pattern. The contribution of the additional predictor variables was non-significant in the ventral tracts ($p > 0.1$). However, letter-based fluency made a significant contribution in the left AF-ft ($p = 0.048$) and category-based fluency made a significant contribution in the right AF-ft ($p = 0.041$). *$p < 0.05$, FDR corrected for 4 clusters. $^{§}p < 0.05$ for the morpheme-based fluency predictor (root or pattern as indicated) within each regression model
